# SALL4 controls cell fate in response to DNA base composition

**DOI:** 10.1101/2020.06.30.179481

**Authors:** Raphaël Pantier, Kashyap Chhatbar, Timo Quante, Konstantina Skourti-Stathaki, Justyna Cholewa-Waclaw, Grace Alston, Beatrice Alexander-Howden, Heng Yang Lee, Atlanta G. Cook, Cornelia G. Spruijt, Michiel Vermeulen, Jim Selfridge, Adrian Bird

## Abstract

Mammalian genomes contain long domains with distinct average compositions of A/T versus G/C base pairs. In a screen for proteins that might interpret base composition by binding to AT-rich motifs, we identified the stem cell factor SALL4 which contains multiple zinc-fingers. Mutation of the domain responsible for AT binding drastically reduced SALL4 genome occupancy and prematurely up-regulated genes in proportion to their AT content. Inactivation of this single AT-binding zinc-finger cluster mimicked defects seen in *Sall4*-null cells, including precocious differentiation of embryonic stem cells and embryonic lethality in mice. In contrast, deletion of two other zinc-finger clusters was phenotypically neutral. Our data indicate that loss of pluripotency is triggered by down-regulation of SALL4, leading to de-repression of a set of AT-rich genes that promotes neuronal differentiation. We conclude that base composition is not merely a passive by-product of genome evolution, but constitutes a signal that aids control of cell fate.

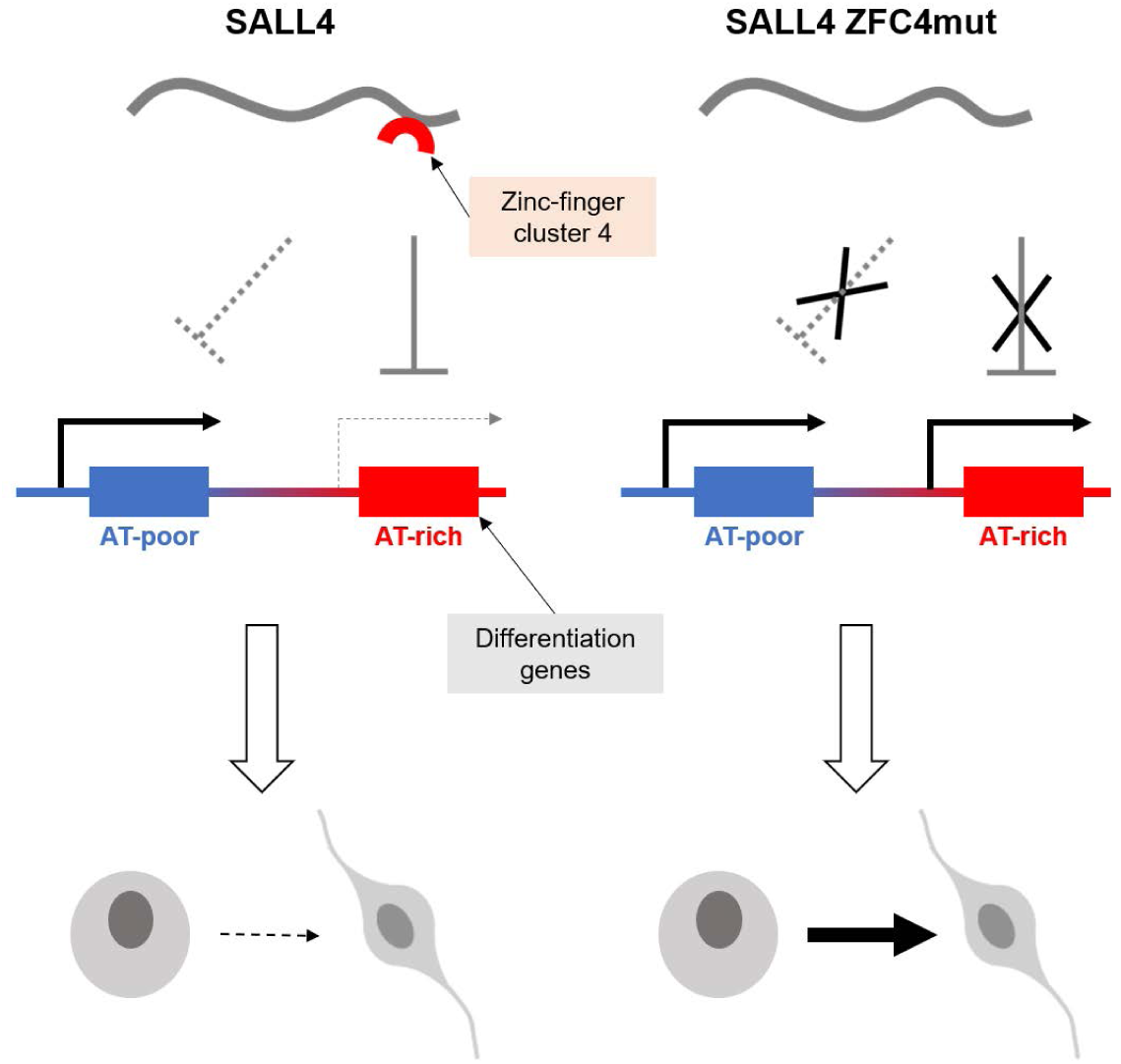

## Introduction

A/T and G/C base pairs are non-randomly distributed within mammalian genomes, forming large and relatively homogenous AT-rich or GC-rich regions that usually encompass several genes together with their intergenic sequences. Base compositional domains are often evolutionarily conserved (Bernardi et al., 1985; Holmquist, 1989; Bickmore and Sumner, 1989; Costantini et al., 2009) and coincide with other genomic features (Bickmore and van Steensel, 2013) including early/late-replicating regions (Hiratani et al., 2009; Holmquist et al., 1982), lamina-associated domains (Meuleman et al., 2013) and topologically associating domains (Jabbari and Bernardi, 2017). Despite these interesting correlations, it is unclear whether conserved AT-rich and GC-rich domains are passive by-products of evolution or whether DNA base composition can play an active biological role (Eyre-Walker and Hurst, 2001; Duret et al., 2006; Arhondakis et al., 2011). Exemplifying this second hypothesis, CpG islands represent conserved GC-rich domains (Bird, 1986) which are specifically bound by proteins recognising unmethylated ‘CG’ dinucleotides (Lee et al., 2001; Voo et al., 2000) to modulate chromatin structure and regulate gene expression (Thomson et al., 2010; Blackledge et al., 2010; Farcas et al., 2012; Wu et al., 2013; He et al., 2013).

Here we tested the hypothesis that AT-rich DNA can be interpreted by specific proteins that recognise short AT-rich motifs whose frequency tracks fluctuations in base composition across the genome (Quante and Bird, 2016). To identify novel AT-binding proteins, we utilized a DNA pulldown-mass spectrometry screen in mouse embryonic stem cells (ESCs) which are pluripotent and can be differentiated in culture. As a top hit we identified SALL4 which is a multi-zinc-finger protein that restrains differentiation of ESCs (Yuri et al., 2009; Miller et al., 2016) and participates in several physiological processes, including neuronal development (Böhm et al., 2008; Sakaki-Yumoto et al., 2006; Tahara et al., 2019), limb formation (Akiyama et al., 2015; Koshiba-Takeuchi et al., 2006) and gametogenesis (Chan et al., 2017; Hobbs et al., 2012; Xu et al., 2017; Yamaguchi et al., 2015). Deletion of the *Sall4* gene leads to embryonic lethality shortly after implantation (Sakaki-Yumoto et al., 2006; Elling et al., 2006; Warren et al., 2007). In humans, failure of SALL4 function is the cause of two severe developmental disorders: the recessive genetic disorder Okihiro syndrome (Al-Baradie et al., 2002; Kohlhase et al., 2002) and embryopathies due to treatment during pregnancy with the drug thalidomide (Donovan et al., 2018; Matyskiela et al., 2018). Despite its biological and biomedical importance, the molecular functions of SALL4 are incompletely understood. The extreme N-terminus interacts specifically with the NuRD co-repressor complex and can account for the transcriptional repression caused by SALL4 recruitment to reporter genes (Lauberth and Rauchman, 2006; Lu et al., 2009). In addition, there is evidence that the zinc-finger clusters bind to DNA (Sakaki-Yumoto et al., 2006; Xiong et al., 2016) or protein partners (Hobbs et al., 2012; Tanimura et al., 2013), though their precise developmental roles are unclear. The present work demonstrates that many of the defects seen in *Sall4*-null ESCs, including precocious differentiation, are mimicked by inactivation of a single zinc-finger cluster that interacts with AT-rich motifs. We go on to show that the ability of SALL4 to sense DNA base composition is essential to restrain transcription of genes that promote differentiation.

## Results

### A screen for AT-binding proteins in embryonic stem cells identifies SALL4

To identify proteins able to sense base composition, we used a DNA affinity purification approach coupled with SILAC-based quantitative mass spectrometry (Spruijt et al., 2013a,b). Mouse ESC protein extracts were mixed with double-stranded DNA probes carrying variable runs of five base pairs composed only of A or T nucleotides (AT-1 and AT-2). As a negative control, matched probes with AT-runs interrupted by G or C nucleotides were used as bait (Ctrl-1 and Ctrl-2). To robustly assess DNA binding specificity, experiments were performed both in the “forward” (heavy-labelled AT-probe *vs* light-labelled Ctrl-probe) and “reverse” (heavy-labelled Ctrl-probe *vs* light-labelled AT-probe) orientations, which were considered as biological replicates. Proteins that bind specifically to AT-runs show a high ratio in the forward experiments (Figure 1A, X-axes) and a low ratio in the reverse experiments (Figure 1A, Y-axes). Mass spectrometry identified a consistent set of AT-binding proteins that largely overlapped between replicate experiments (Figure S1A) and between unrelated AT-rich probes (Figure S1B). High confidence hits included proteins with well characterised AT-binding domains such as AT-hooks (Aravind and Landsman, 1998; Filarsky et al., 2015) (HMGA1, HMGA2, PRR12, BAZ2A) and “AT-rich interaction domains” (Patsialou et al., 2005) (ARID3A, ARID3B, ARID5B), thereby validating the screen (Figure 1A and Table S1). Three Spalt-like (SALL) family proteins (Sweetman and Münsterberg, 2006) (SALL1, SALL3, SALL4) and most components of the NuRD complex (Tong et al., 1998; Wade et al., 1998; Xue et al., 1998; Zhang et al., 1998) were also recovered (Figure 1A). The most consistently enriched protein in our mass spectrometry screen was SALL4, whose AT-binding we confirmed by Western blot analysis using a variety of probes with one (AT-3) or more AT-runs (Figure S1C). Considering their reported interaction with NuRD (Lauberth and Rauchman, 2006), we suspected that SALL proteins might be responsible for recruitment of this co-repressor complex to AT-rich DNA. To test this, we used extracts from mouse ESCs in which the *Sall4* gene is disrupted (*S4KO* ESCs) (Miller et al., 2016; Sakaki-Yumoto et al., 2006). As predicted, recovery of NuRD components by AT-rich DNA was greatly reduced compared to *wild-type* (*WT*) in the absence of SALL4 (Figure 1B).

**Figure 1:**
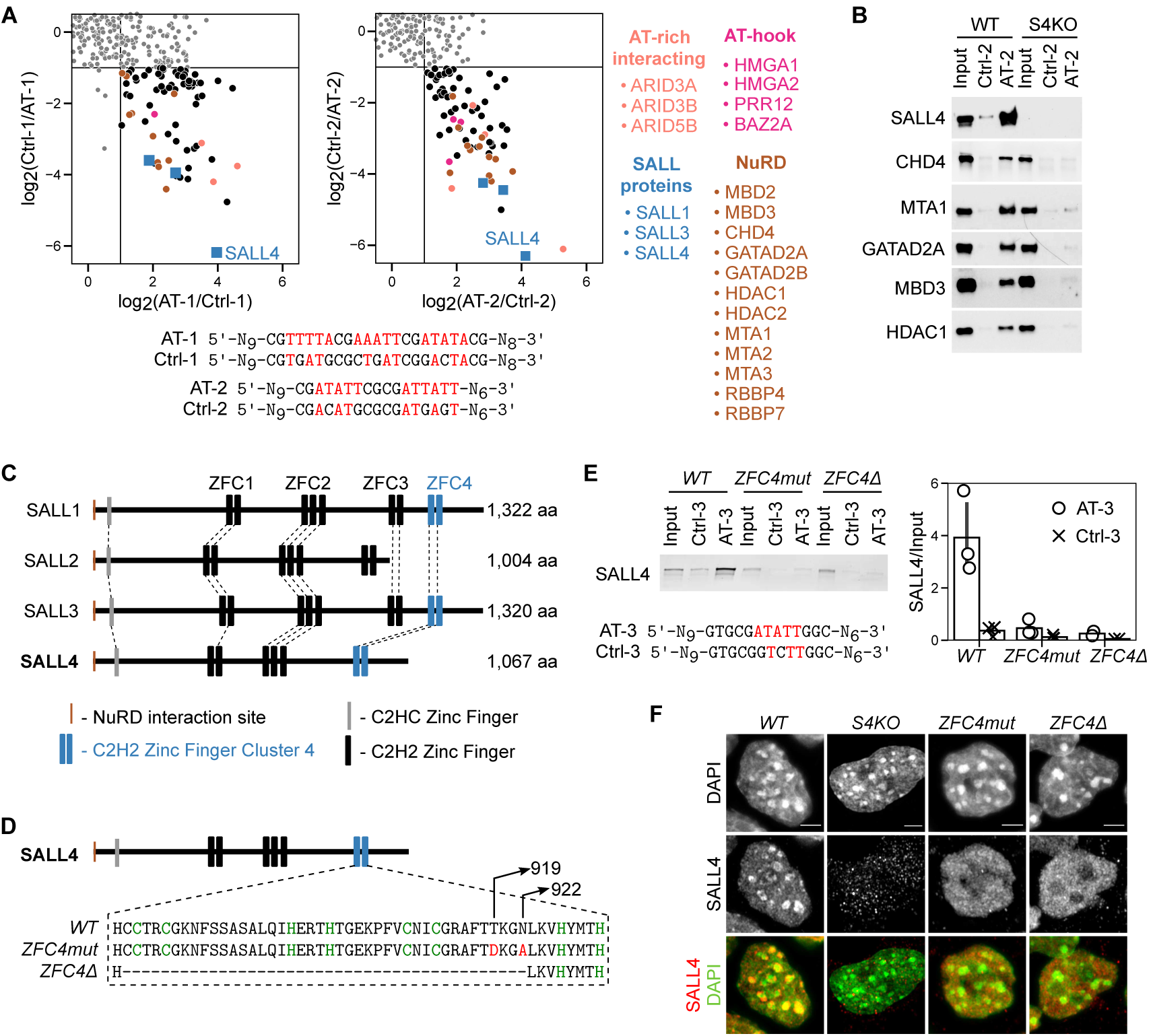
Identification of novel AT-binding proteins in embryonic stem cells by DNA pulldown-mass spectrometry. **A.** Scatter plots representing SILAC-based DNA affinity purifications, comparing AT-rich DNA probes (AT-1, AT-2) with control probes having interrupted AT-runs (Ctrl-1, Ctrl-2). The ratio of the quantified proteins in the forward experiment (Heavy-AT/Light-Ctrl) is plotted on the X-axis and the ratio for the same proteins in the reverse experiment (Heavy-Ctrl/Light-AT) is plotted on the Y-axis. Proteins were considered to be specific interactors when showing at least a 2-fold ratio in both the forward and reverse experiments. These proteins cluster in the bottom right quadrant. **B.** DNA pulldown with AT-rich (AT-2) or control (Ctrl-2) probes followed by Western blot analysis for SALL4 and NuRD components using *wild-type* (*WT*) or *Sall4* knockout (*S4KO*) ESC protein extracts. **C.** Protein alignment of mouse SALL family members indicating conserved protein domains, including C2H2 zinc-finger clusters (ZFC1-4). **D.** Diagram showing the mutations or deletion introduced within SALL4 ZFC4 by CRISPR/Cas9. **E.** DNA pulldown with AT-rich (AT-3) or control (Ctrl-3) probe followed by Western blot analysis for SALL4 using *WT* or *Sall4 ZFC4mut/*∆ ESC protein extracts. SALL4 levels were quantified and normalised to input. Data points indicate independent replicate experiments, and error bars standard deviation. **F.** SALL4 immunofluorescence in the indicated ESC lines. DNA was stained with DAPI, showing dense clusters of AT-rich pericentric chromatin. Scale bars: 3µm.

### SALL4 binds to short AT-rich motifs via C2H2 zinc-finger cluster 4 (ZFC4)

Mammalian genomes encode four SALL family proteins (SALL1-4) which each contain several clusters of C2H2 zinc-fingers. Based on similarities in amino acid sequence between family members, the clusters are classified as ZFC1-4 (Figure 1C). SALL1, SALL3 and SALL4 all possess ZFC4 (Figure S1D), but SALL2 lacks this domain and was not recovered in our screen for AT-binding proteins. ZFC4 of both SALL1 and SALL4 was previously shown to interact with AT-rich heterochromatin in transfection assays (Sakaki-Yumoto et al., 2006; Yamashita et al., 2007), suggesting that it might be responsible for AT-binding. To further characterise this domain, we used CRISPR/Cas9 to either delete ZFC4 (*ZFC4*∆) or mutate two residues (T919D, N922A; mutated residues shown in red) that we predicted would be involved in DNA binding (*ZFC4mut*) (Figure 1D). Homozygous mouse ESC lines expressing both mutated SALL4 proteins were obtained (Figure S1E), both of which retained the ability to interact with NuRD components by co-immunoprecipitation (Figure S1F). The interaction of SALL4 ZFC4mut or ZFC4∆ proteins with AT-rich sequences was drastically reduced (>10-fold) by inactivation of ZFC4, as assessed by the DNA pulldown assay (Figure 1E). This strongly suggests that the ZFC4 domain of SALL4 is primarily responsible for pulldown by AT-rich DNA. We next explored the *in vivo* DNA binding properties of SALL4 ZFC4 in our mutant ESC lines. Heterochromatic foci, identified by DAPI staining in mouse cells, contain a high concentration of AT-rich satellite DNA (Matsuda and Chapman, 1991; Cerda et al., 1999; Guenatri et al., 2004), and therefore provide a low-resolution cellular assay for preferential AT-binding. Immunostaining with a SALL4 antibody recognising a preserved epitope in the two mutant proteins revealed a striking loss of ZFC4mut and ZFC4∆ protein localisation at DAPI-dense foci (Figure 1F), further confirming that this zinc-finger cluster is necessary for AT targeting.

To define the sequence preference of the purified ZFC4 domain, we performed HT-SELEX whereby a library of initially random DNA sequences immobilised on beads was subjected to repeated cycles of binding and amplification (Jolma et al., 2010; Nitta et al., 2015). After 0, 1, 3 or 6 cycles, DNA was analysed by high-throughput sequencing to detect enriched motifs. For comparison, we performed HT-SELEX on other SALL4 zinc-finger clusters ZFC1 and ZFC2 (Figure 2A), and also included a sample without added proteins to control for PCR bias. Strikingly, with increasing cycles of ZFC4 binding, the base composition of the whole library gradually shifted towards higher AT-content, but this effect was not seen with ZFC1, ZFC2 or the negative control (Figure 2B). Progressive A/T motif enrichment was also apparent for ZFC4 alone (Figure 2C). To determine the minimum number of A/T required for enrichment, we separated the oligomers (k-mers) into different groups (Figure 2C). Enrichment was only detectable in k-mers containing 4 or more A/T base pairs, suggesting that this is the minimum target sequence. After 6 cycles, the most enriched SELEX motif was ‘ATATT’ (Figure S2A), which also corresponds to the preferred sequence identified by DNA pulldown using all possible combinations of AT 5 mers (Figure S2B). However, this is one of several target sequences, as multiple other AT-rich sequences had similar enrichment scores (Figure 2B, S2A). All 16 possible A/T 5-mers are enriched more than G/C containing 5-mers, with k-mers containing TATA among the least favoured AT-rich motifs for ZFC4 binding. We conclude that ZFC4 targets a broad range of short motifs that are composed only of A and T, whereas ZFC1 and ZFC2 showed no apparent DNA sequence specificity.

**Figure 2:**
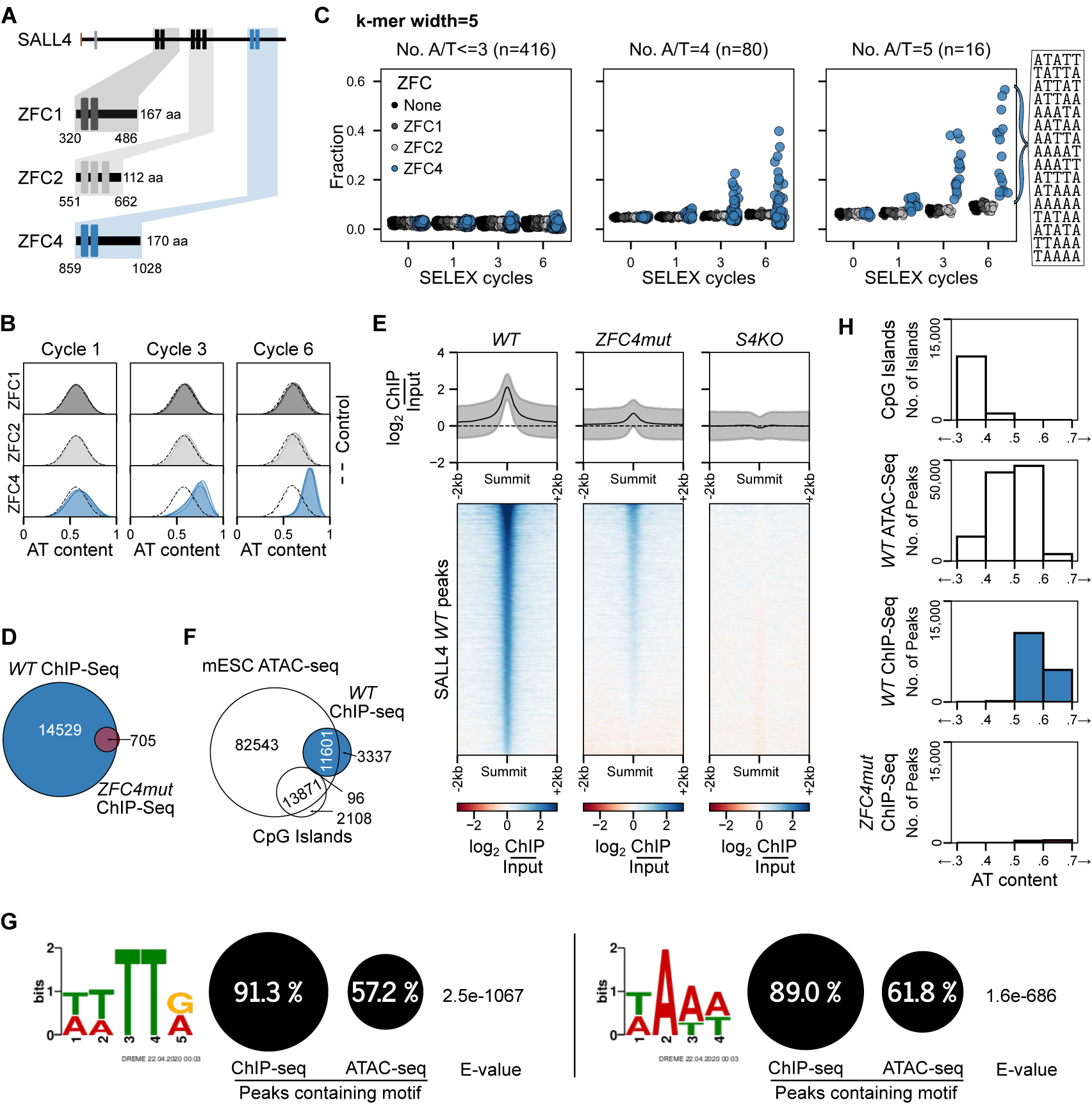
Characterisation of SALL4 C2H2 zinc-finger cluster 4 (ZFC4) DNA binding *in vitro* and *in vivo*. **A.** SALL4 ZFC1, ZFC2 and ZFC4 protein fragments used for *in vitro* HT-SELEX experiments. A sample without added protein was used as a negative control. **B.** Base composition of HT-SELEX libraries following 1, 3 and 6 cycles with ZFC1 (dark grey), ZFC2 (light grey) and ZFC4 purified proteins (blue). **C.** Relative enrichment of 5-mer motifs categorised by AT-content at cycle 0, 1, 3 and 6 of HT-SELEX with SALL4 ZFC1 (dark grey), ZFC2 (light grey), ZFC4 (blue) and negative control (black) samples. All 16 possible A/T 5-mer motifs are ordered according to their enrichment at cycle 6 of ZFC4 HT-SELEX. **D.** Venn diagram showing the overlap of SALL4 ChIP-seq peaks between *WT* and *ZFC4mut* ESCs. **E.** Profile plot and heatmap showing SALL4 ChIP-seq signal at SALL4 *WT* ChIP-seq peaks in the indicated cell lines. **F.** Venn diagram showing the overlap of SALL4 ChIP-seq peaks detected in *WT* ESCs with ATAC-seq peaks (accessible chromatin) and CpG islands. **G.** Results from *de novo* motif analysis at SALL4 *WT* ChIP-seq peaks (summit +/− 250bp) showing the relative frequency of each DNA motif and its associated E-value. ATAC-seq peaks were used as a control for regions of accessible chromatin. **H.** Analysis of the DNA base composition surrounding SALL4 ChIP-seq peaks (summit +/− 250bp) in *WT* (blue) and *ZFC4mut* (red) ESCs. CpG islands and ATAC-seq peaks coincide with regions of accessible chromatin and are shown for comparison.

### ZFC4 mutation drastically reduces SALL4 chromatin binding *in vivo*

To assess the influence of ZFC4 on SALL4 chromatin occupancy *in vivo*, we performed ChIP-seq using two anti-SALL4 antibodies (one monoclonal, one polyclonal) recognising a C-terminal epitope which is distant from C2H2 zinc-finger clusters. We first determined antibody specificity (Kidder et al., 2011; Landt et al., 2012; Uhlen et al., 2016) by assessing SALL4 ChIP signal in *S4KO* ESCs as a negative control. Over 15,000 non-specific ChIP-seq peaks were observed with the polyclonal anti-SALL4 antibody, compared with only 280 peaks for the monoclonal antibody (Figure S2C, S2D). We therefore analysed exclusively data obtained with the anti-SALL4 monoclonal antibody, considering only ChIP-seq peaks that were consistent between independent replicate experiments in *WT* or *ZFC4mut* ESCs (Figure S2E). In agreement with its reported localisation at enhancers (Miller et al., 2016; Xiong et al., 2016), we observed that SALL4 ChIP-seq peaks in *WT* cells were enriched in the histone marks H3K27ac and H3K4me1 (Chronis et al., 2017), which typically mark these genomic sites (Figure S2F). Strikingly, *ZFC4mut* cells lost ~95% of ChIP-seq peaks compared to *WT* (Figure 2D). Heatmaps confirmed the depletion of SALL4 peaks, although we noted a small amount of bound ZFC4mut at a subset of *WT* binding sites (Figure 2E).

We compared wildtype SALL4 binding sites as a whole with regions of open chromatin identified by ATAC-seq, which detects accessible DNA, including enhancers and promoters. SALL4 peaks largely coincide with a subset of ATAC-seq peaks, while avoiding CpG island promoters (Figure 2F). The AT-binding specificity of SALL4 suggests that this protein might preferentially associate with open chromatin sites that are more AT-rich than average. The complete absence of SALL4 at ATAC-seq peaks within CpG islands (Figure 2F), within which runs of As and Ts are rare, is compatible with this notion. To quantify this effect, we used *de novo* motif analysis to determine whether SALL4 peaks were consistent with a bias towards AT-rich motifs. Firstly, by seeking recurrent motifs (<8 base pairs) coincident with SALL4 peaks we identified short AT-rich motifs that were highly enriched at the majority (~90%) of SALL4 binding sites compared with lower levels of enrichment (~60%) in open chromatin generally (Figure 2G). As a second approach, we determined the base composition at SALL4-bound regions by analysing the DNA sequences surrounding SALL4 ChIP-seq peak summits (+/− 250bp). SALL4 binding sites are relatively AT-rich (50-70% AT) (Figure 2H) compared with ATAC-seq peaks as a whole (40-60% AT) (Figure 2H). Taken together, the data suggest that AT-motif binding is responsible for the presence of SALL4 at a subset of open chromatin sites.

### SALL4 ZFC4 represses the expression of early differentiation genes in a base composition-dependent manner

To determine whether SALL4 binding to AT-rich DNA causes gene expression changes that correlate with base composition, we performed RNA-seq in *WT*, *ZFC4mut*, *ZFC4*∆ and *S4KO* ESCs. *Sall4* gene knockout resulted in the dysregulation of several thousand genes (Figure 3A). Both ZFC4 mutations caused the dysregulation of fewer genes, many of which overlapped with those affected in *S4KO* cells (Figure 3A). To test the relationship between AT composition and gene expression, genes differentially regulated in both *ZFC4mut* and *ZFC4*∆ ESCs (Figure 3A, red filling) were divided into five equal categories according to AT-content across the entire transcription unit (Figure 3B), and the level and direction of transcriptional change was compared between them. In agreement with our hypothesis, genes differentially regulated in *ZFC4mut/*∆ cells showed progressively increased up-regulation with rising AT-content (Figure 3C). To quantify the strength of the relationship between AT-content and gene expression, we fitted a linear regression model and calculated coefficient estimates. This independent approach, which reveals the variation in gene expression that can be attributed to base composition, confirmed that the positive relationship between AT-content and up-regulation in the ZFC4 mutants is significant (FDR<0.01; see Methods and Table S2; Figure S3A). In contrast, genes differentially regulated in *S4KO*, but not in either of the *ZFC4 mutant* ESCs (Figure 3A, grey filling), showed a non-significant correlation (FDR>0.01) and an effect size close to zero (Figure 3D, S3B and S3C). The results show that the subset of SALL4-regulated genes that is dependent on ZFC4 is repressed in pluripotent cells according to the AT-richness of their genomic setting.

**Figure 3:**
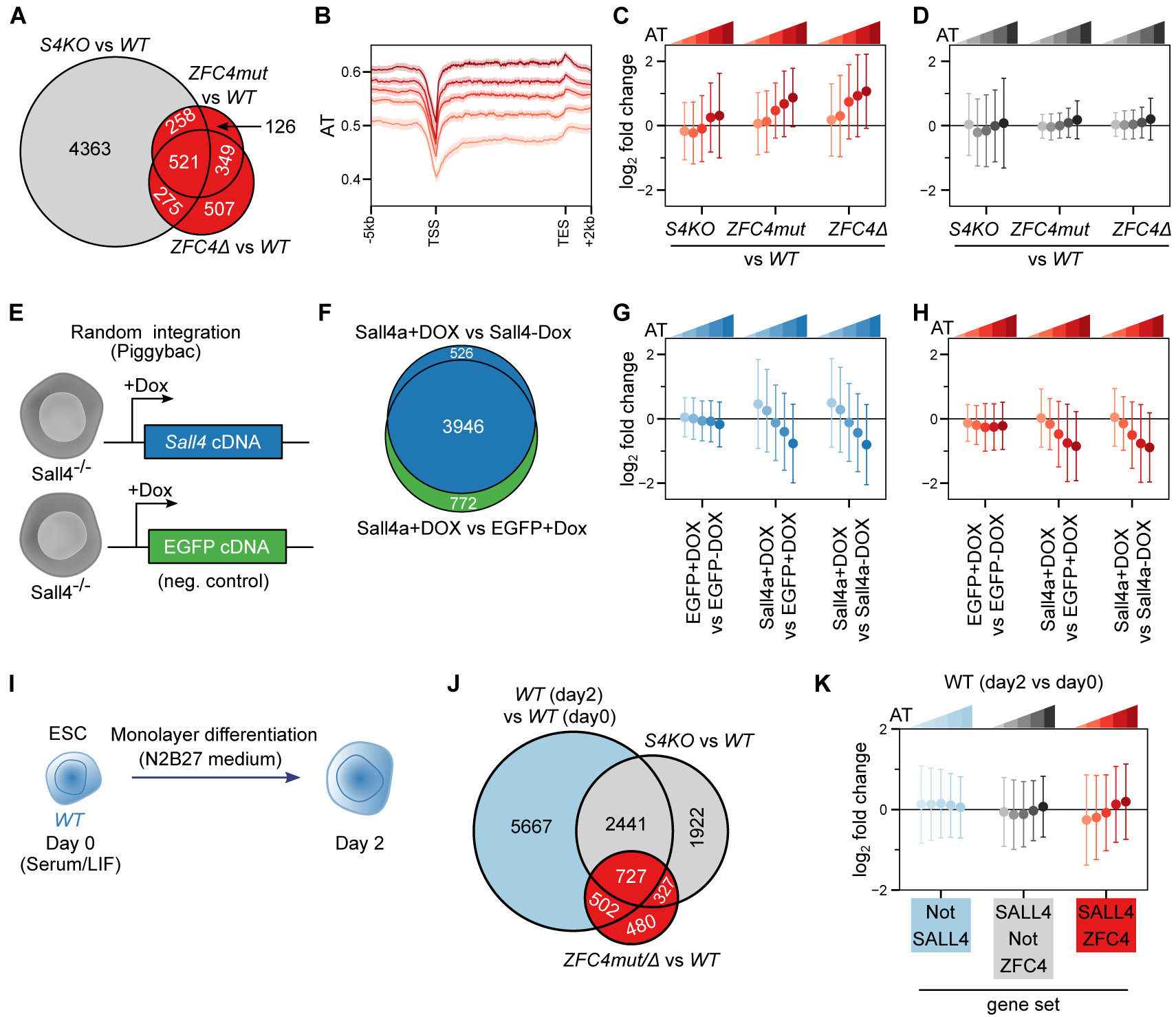
SALL4-mediated transcriptional regulation in relation to DNA base composition. **A.** Venn diagram showing the overlap of differentially expressed genes detected by RNA-seq between *S4KO*, *ZFC4mut* and *ZFC4*∆ ESCs. ZFC4-regulated genes are indicated in red, and ZFC4-independent genes in grey. **B.** Profile plot showing the density of A/T nucleotides around the transcription unit of ZFC4-regulated genes divided into five equal categories according to AT-content. TSS: Transcription start site, TES: Transcription end site. **C, D.** Correlation between gene mis-regulation (log_2_ fold-change *vs WT*) and DNA base composition in *Sall4* mutant ESCs. ZFC4-regulated (C) and ZFC4-independent (D) genes were divided into five equal categories depending on their AT-content. **E.** Diagram representing *Sall4* knockout ESC lines carrying SALL4 or EGFP (control) expression constructs under control of a doxycycline-inducible promoter. **F.** Venn diagram showing the overlap of differentially expressed genes detected by RNA-seq following a 48h doxycycline induction in the ESC lines presented in panel E. SALL4-responsive genes are indicated in blue, and EGFP-responsive genes in green. **G, H.** Correlation between SALL4-induced gene expression changes and DNA base composition. SALL4-responsive (G) and ZFC4-regulated (H) genes were divided into five equal categories depending on their AT-content, and their relative expression levels were analysed in the indicated ESC lines. **I.** Diagram showing the protocol used to characterise early differentiation of *WT* ESCs. **J.** Venn diagram showing the overlap between genes changing during early differentiation of *WT* cells (day 0 *vs* day 2) with genes de-regulated in *Sall4* mutant ESCs. Genes were divided into three categories: SALL4-independent genes (light blue), SALL4-dependent genes controlled by ZFC4 (red) and SALL4-dependent genes not controlled by ZFC4 (grey). **K.** Correlation between gene expression changes occurring during early differentiation and DNA base composition in *WT* cells. SALL4-independent genes (light blue), SALL4-dependent genes controlled by ZFC4 (red) and SALL4-dependent genes not controlled by ZFC4 (grey) were divided into five equal categories depending on their AT-content, and their relative expression levels were analysed at day 2 of differentiation.

To further test the hypothesis that AT-binding by ZFC4 mediates repression according to base composition, we examined the reverse situation of SALL4 over-expression on transcription. This was performed by expressing SALL4, or as a negative control EGFP, from a doxycycline-inducible promoter following random integration of expression constructs in *S4KO* ESCs (Figure 3E). After 48 hours of induction, SALL4 was robustly over-expressed in these cells (Figure S3D, S3E). To characterise the effect of SALL4 re-expression on transcription, we performed RNA-seq on induced (+Dox) and non-induced (-Dox) cell lines (Figure 3F). As expected, gene expression changes in cells over-expressing SALL4 were anti-correlated with expression changes seen in *S4KO* cells (Figure S3F). Separation of differentially expressed genes into categories according to their AT-content as before revealed that SALL4 expression caused transcriptional repression that was strikingly proportional to the base composition of the affected genes (Figure 3G and S3G). A similar relationship was observed when looking at genes differentially regulated in *ZFC4mut/*∆ ESCs (Figure 3H). Linear regression analysis again confirmed the significance of these relationships (Figure S3H and S3I). As a control, we applied the same analysis to the minority of genes whose expression was altered in response to EGFP-induction (Figure 3F, green filling). In this case, there was no apparent relationship between fold-change and base composition (Figure S3J and S3K), as confirmed by quantitative analysis (Figure S3L). Together our results strongly suggest that SALL4 directly regulates gene expression in response to base composition via its zinc-finger cluster ZFC4.

Interestingly, gene ontology (GO) analysis of ZFC4-regulated genes identified GO terms associated with neuronal differentiation, morphogenesis, gonad development and kidney development (Table S3), all of which are adversely affected in *Sall4* knockout mice and embryos (Böhm et al., 2008; Sakaki-Yumoto et al., 2006; Tahara et al., 2019; Akiyama et al., 2015; Koshiba-Takeuchi et al., 2006; Chan et al., 2017; Hobbs et al., 2012; Xu et al., 2017; Yamaguchi et al., 2015; Warren et al., 2007). This suggests the possibility that SALL4 plays an essential role in the transition between self-renewing ESCs and the differentiated state by preferentially suppressing the expression of AT-rich developmental genes, thus preventing premature loss of pluripotency. If so, AT-rich genes that are aberrantly up-regulated in the absence of ZFC4 should be activated during the normal differentiation programme of *WT* cells (Rao et al., 2010). To test this, we performed RNA-seq on *WT* ESCs following two days of monolayer differentiation (Figure 3I) (Aubert et al., 2002). Although they represent a small fraction of all transcriptional changes taking place at these stages, SALL4-regulated genes overlapped significantly with genes whose expression changes naturally between day 0 (ESCs) and day 2 of differentiation (Figure 3J). Importantly, ZFC4-regulated genes, but not other categories of genes, are up-regulated at this early stage in proportion to AT-richness (Figure 3K, S3M and S3N). Thus, AT-rich genes that are repressed by SALL4 in ESCs are activated soon after the exit from pluripotency.

### SALL4 ZFC4 is critical for neuronal differentiation and embryonic development

Previous work demonstrated that disruption of the *Sall4* gene leads to increased stem cell differentiation (Yuri et al., 2009; Miller et al., 2016). To test whether disrupting ZFC4 alone leads to phenotypic defects, we compared *ZFC4mut* and *S4KO* ESCs. Consistent with previous evidence showing that SALL4 is dispensable for the maintenance of pluripotency (Yuri et al., 2009; Sakaki-Yumoto et al., 2006; Tsubooka et al., 2009), both *S4KO* and *ZFC4mut* ESCs expressed normal levels of OCT4 (Figure S4A) and showed efficient self-renewal, with a modest decrease observed in *S4KO* ESCs (Figure S4B). Next, we used a monolayer differentiation assay, as described above, to assess the propensity of these cell lines to acquire a neuronal fate. After 5 days in N2B27 medium, ESCs lacking SALL4 or expressing a ZFC4 mutant protein generated many more TUJ1-positive cells compared to *WT* cells (Figure 4A). Further confirming increased neuronal differentiation, RT-qPCR analyses identified increased transcription of *Tuj1* (4-12 fold), *Ascl1* (3-6 fold) and *Nestin* (~2 fold) in *S4KO*, *ZFC4mut* and *ZFC4*∆ ESCs at day 5 of differentiation (Figure 4B). By this assay, inactivation of ZFC4 phenocopies complete loss of SALL4 protein.

**Figure 4:**
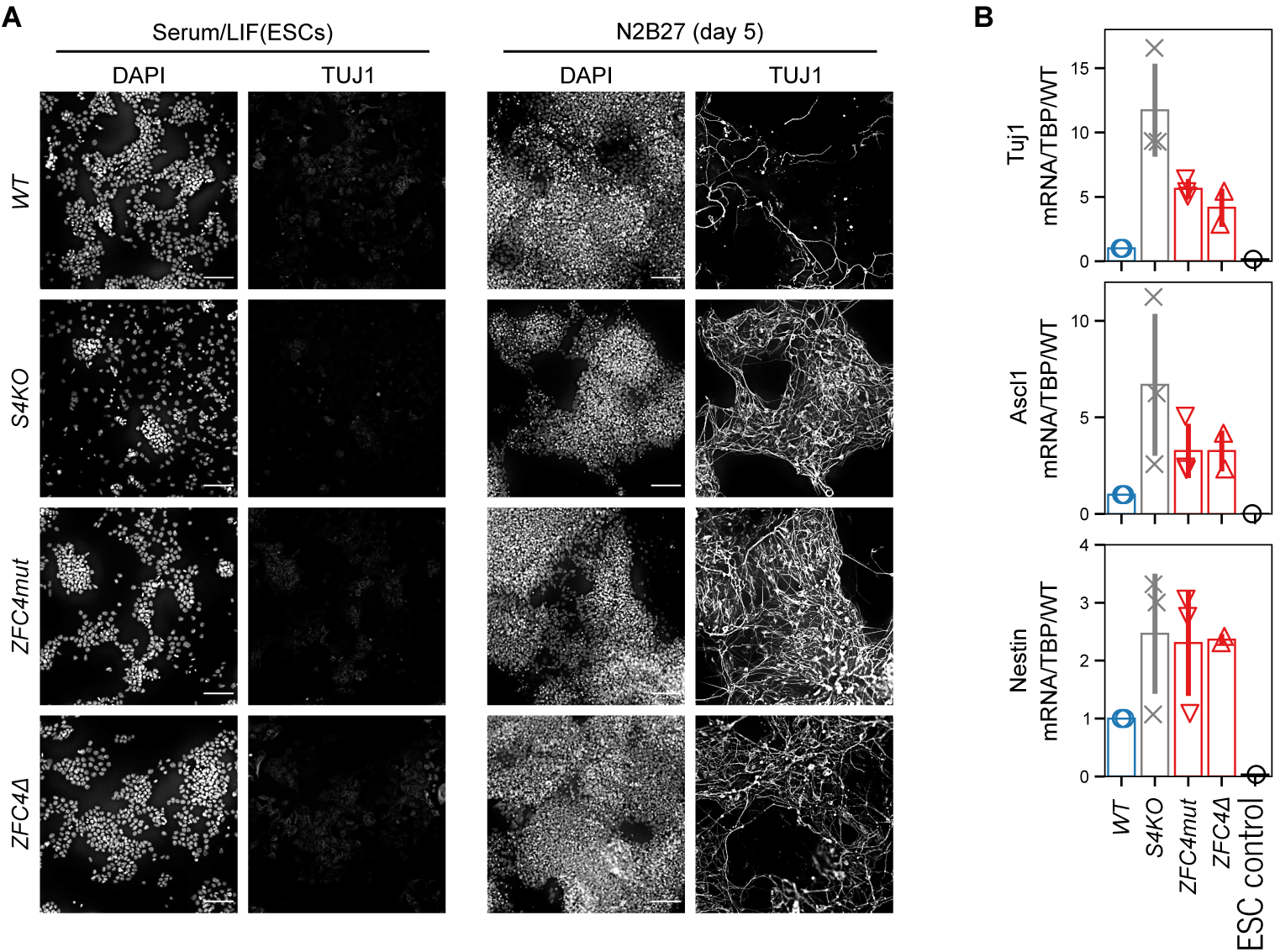
Phenotypic characterisation of SALL4 ZFC4 mutation during neuronal differentiation. **A.** TUJ1 immunofluorescence in the indicated ESC lines cultured in serum/LIF medium, and following differentiation for 5 days in N2B27 medium. DNA was stained with DAPI. Scale bars: 100µm. **B.** RT-qPCR analysis of the neuronal markers Tuj1, Ascl1 and Nestin in the indicated cell lines following differentiation for 5 days in N2B27 medium. Transcripts levels were normalised to TBP and expressed relative to *WT*. Data points indicate independent replicate experiments and error bars standard deviation.

In order to observe the effects of ZFC4 mutation on embryonic development, we generated a *ZFC4mut* mouse line by blastocyst injection of heterozygous *Sall4*^*ZFC4mut/WT*^ ESCs. F1 mice were crossed and their progeny analysed at different stages of development. While *ZFC4mut* homozygous embryos were obtained at Mendelian ratios during early development, none survived until birth (Figure 5A and 5B). By E10.5, homozygous embryos presented gross morphological abnormalities, which were not observed in controls (Figure 5C). Importantly, the ZFC4mut protein was expressed at levels similar to those seen in *WT* embryos (Figure S5A and S5B). Early embryonic mortality of *ZFC4* mutant mice is reminiscent of the phenotype observed in *Sall4* knockout mice, although the latter die earlier in development, shortly after implantation (by E5.5-6.5) (Sakaki-Yumoto et al., 2006; Elling et al., 2006). Taken together, our *in vitro* and *in vivo* experiments indicate that mutation of ZFC4 alone phenocopies important aspects of the *Sall4* knockout phenotypes seen in both ESCs and embryos. It follows that this DNA binding domain is a key contributor to SALL4 biological function.

**Figure 5:**
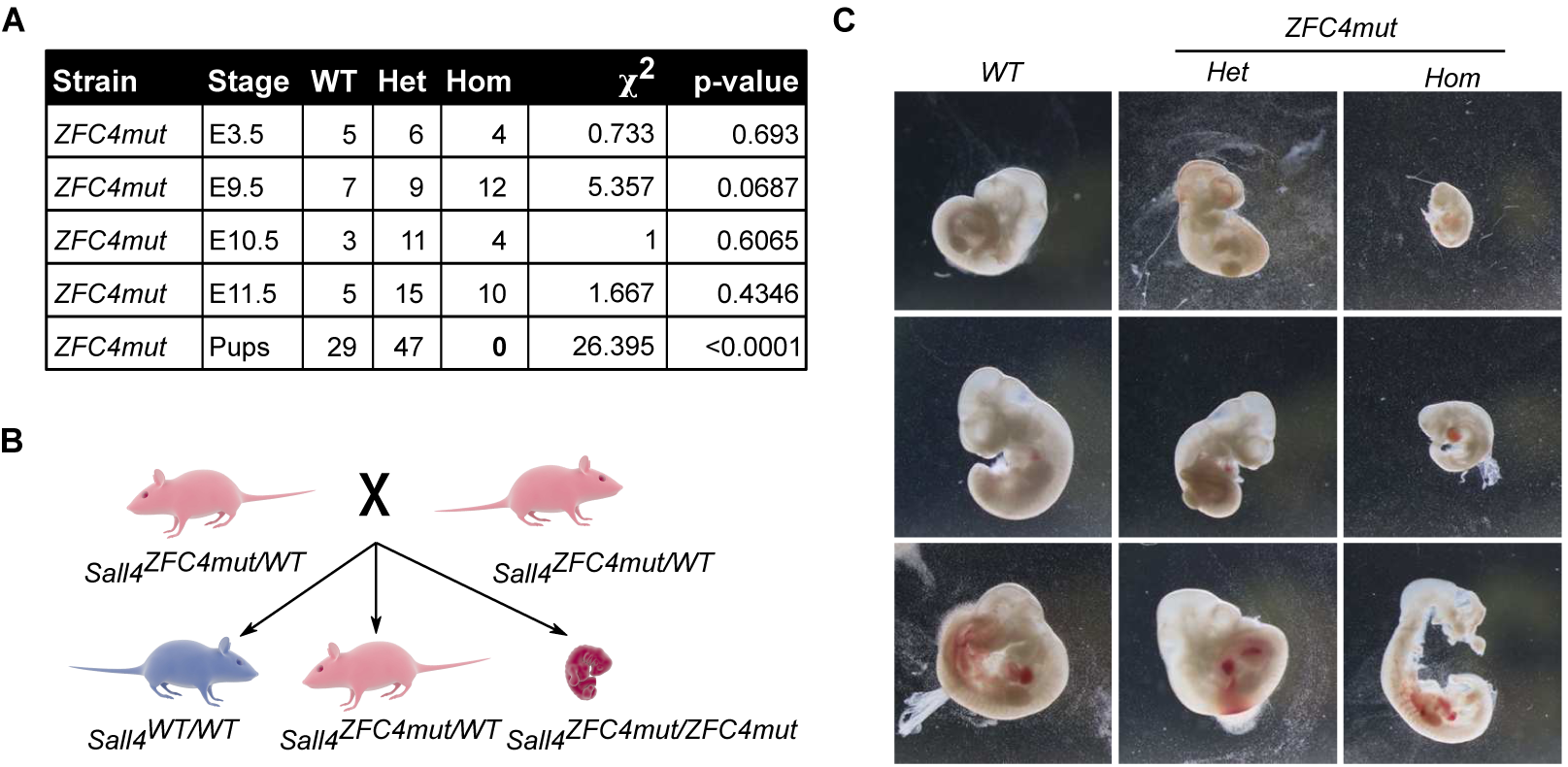
Phenotypic characterisation of SALL4 ZFC4 mutation during embryonic development. **A.** Table showing the number of live pups and embryos at different stages of development, and their associated genotype. Animals were crossed to obtain *ZFC4mut* heterozygous (*Het*), homozygous (*Hom*), or *WT* progeny. **B.** Diagram showing the results from crossing *ZFC4mut* heterozygote mice. *ZFC4mut* homozygous animals die during embryonic development. **C.** Representative images of *WT*, *ZFC4mut* heterozygous (*Het*) and homozygous (*Hom*) embryos at E10.5, taken at the same magnification.

### C2H2 zinc-finger clusters 1 and 2 are dispensable for SALL4 function in ESCs

SALL4 contains two C2H2 zinc-finger clusters, ZFC1 and ZFC2, in addition to ZFC4. To determine their contribution to SALL4 function, we used CRISPR/Cas9 to delete the central segment of endogenous SALL4 protein which contains zinc-finger clusters ZFC1 and ZFC2, while leaving ZFC4 and the N-terminal domain intact (Figure 6A). ESCs homozygous for this *ZFC1-2*∆ knock-in allele lack full length SALL4, but, as expected, ZFC1-2∆ protein retained the ability to interact with SALL1 and NuRD components (Figure S6A). Immunostaining showed that ZFC1-2∆ resembled *WT* SALL4 by being enriched at heterochromatic foci, indicating that ZFC4 binding to this AT-rich DNA *in vivo* is unaffected by the internal deletion (Figure 6B). To characterise ZFC1-2∆ chromatin binding in more detail, we performed ChIP-seq (Figure S6B), as described above. In contrast to the dramatic effect of ZFC4 inactivation on SALL4 ChIP-seq peaks, ZFC1-2∆ occupancy of the genome closely resembled that of *WT* SALL4 (Figure S6C). In addition, both the average ChIP-seq signal (Figure 6C) and AT-rich profile (Figure 6D) of *WT* SALL4 peaks were preserved in *ZFC1-2*∆ cells. We conclude that ZFC1 and ZFC2 contribute minimally to the genome binding profile of SALL4, further supporting the view that ZFC4 is the primary determinant of DNA binding. Comparative RNA-seq analysis between *WT*, *ZFC4mut* and *ZFC1-2*∆ ESCs revealed that SALL4 ZFC1-2∆ and ZFC4mut affect largely non-overlapping sets of genes (Figure 6E). The effects of ZFC1-2∆ on transcription were independent of base composition, whereas ZFC4 regulated genes in proportion to their AT-richness (Figure 6F, S6D, S6E and S6F). Finally, we examined the phenotypic consequences of ZFC1-2 deletion by assaying monolayer neuronal differentiation of our mutant ESCs. Unlike *S4KO* and *ZFC4mut* ESCs, *ZFC1-2*∆ cells did not show evidence of increased differentiation as assessed by TUJ1 immunofluorescence (Figure 6G) and RT-qPCR analysis of neuronal markers at day 5 of differentiation (Figure S6G).

**Figure 6:**
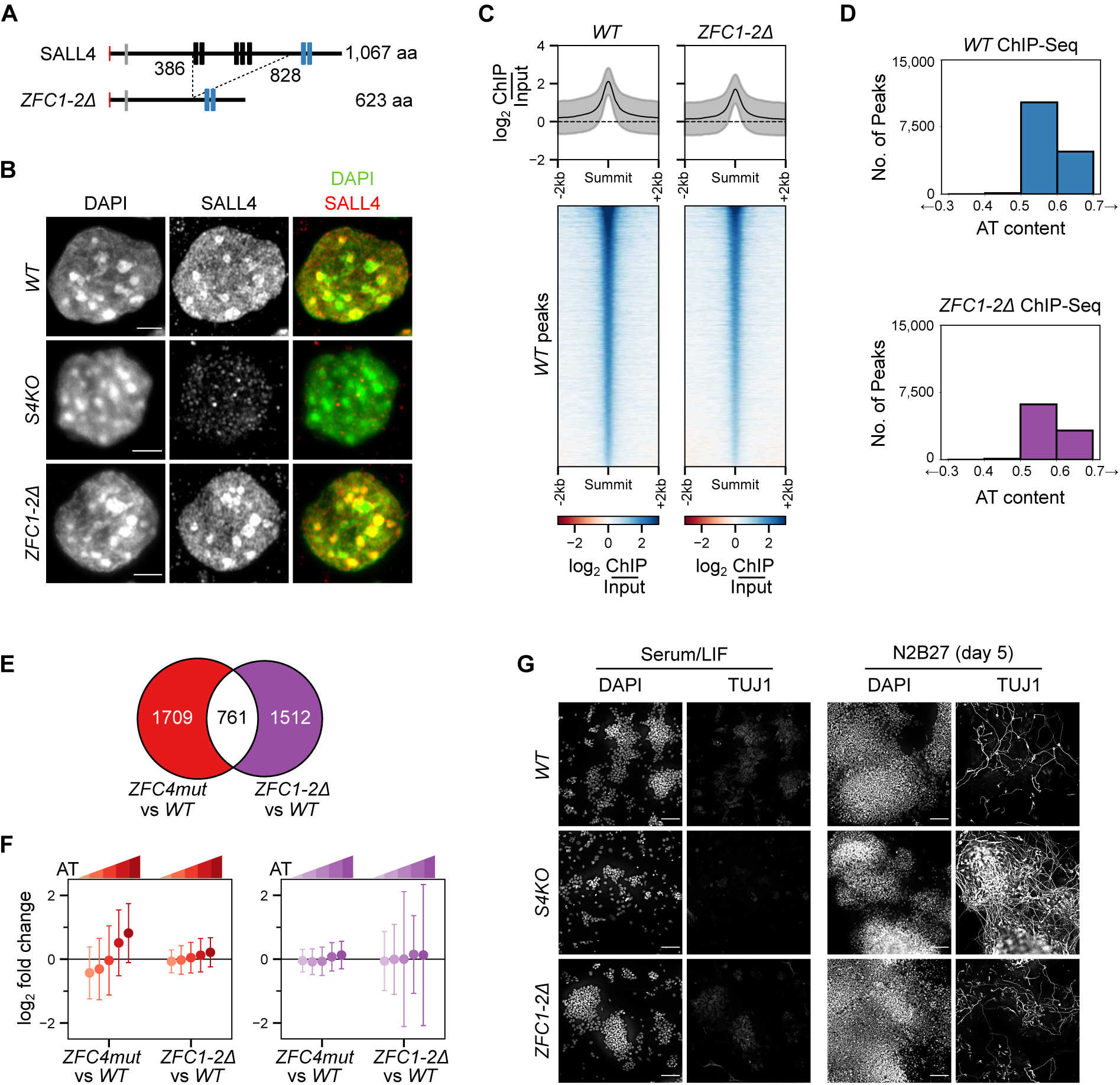
Characterisation of SALL4 C2H2 zinc-finger clusters 1 and 2 in ESCs. **A.** Diagram showing the in frame deletion of SALL4 within the *Sall4* coding sequence, generated by CRISPR/Cas9. **B.** SALL4 ZFC1-2∆ localisation determined by immunofluorescence in the indicated ESC lines. DNA was stained with DAPI, showing dense clusters of AT-rich pericentric chromatin. Scale bars: 3µm. **C.** Heatmap and profile plot showing SALL4 ChIP-seq signal at SALL4 *WT* ChIP-seq peaks in the indicated cell lines. **D.** Analysis of the DNA base composition surrounding SALL4 ChIP-seq peaks (summit +/− 250bp) in *WT* (blue) and *ZFC1-2*∆ (purple) ESCs. **E.** Venn diagram showing the overlap of differentially expressed genes detected by RNA-seq between *ZFC4mut* and *ZFC1-2*∆ ESCs. ZFC4-regulated genes are indicated in red and ZFC1/2-regulated genes in purple. **F.** Correlation between gene mis-regulation (log_2_ fold-change *vs WT*) and DNA base composition in *Sall4* mutant ESCs. ZFC4-regulated (red) and ZFC1/2-regulated (purple) genes were divided into five equal categories depending on their AT-content. **G.** TUJ1 immunofluorescence in the indicated ESC lines cultured in serum/LIF medium, and following differentiation for 5 days in N2B27 medium. DNA was stained with DAPI. Scale bars: 100µm.

To further characterise the differentiation defects of *Sall4* mutant ESCs, we performed an RNA-seq time-course experiment with *WT*, *S4KO*, *ZFC4mut* and *ZFC1-2*∆ cell lines at day 0 (ESCs), day 2 and day 5 of the differentiation protocol (Figure 7A). In agreement with our previous base composition analyses, absence of SALL4 or inactivation of ZFC4 weakened repression, leading to premature activation of ZFC4-regulated AT-rich genes at all differentiation time points (Figure S7A). In contrast, ZFC1/2 regulated genes showed no preferential up-regulation during differentiation, and no correlation with base composition in any of the cell lines (Figure S7B). Moreover, PCA analysis showed that *WT* and *ZFC1-2*∆ samples clustered together at all time points, while *S4KO* and *ZFC4mut* formed an independent cluster at days 2 and 5 (Figure S7C). Accordingly, differential expression analysis across our time series revealed few differences between *WT* and *ZFC1-2*∆, while the transcriptomes of *S4KO* and *ZFC4mut* were significantly disturbed (Figure 7B). Emphasising the similarity of *S4KO* and *ZFC4mut*, genes differentially regulated in these cell lines were highly correlated both at day 2 and 5 of differentiation (Figure 7C and S7D). Also, genes associated with neuronal differentiation were up-regulated in both cell lines, whereas expression of these genes in *ZFC1-2*∆ cells was unaffected (Figure 7D). We conclude that the characteristic premature differentiation phenotype associated with SALL4 deficiency is mimicked by inactivation of ZFC4, but not by a large deletion of the central domain that includes ZFC1 and ZFC2.

**Figure 7:**
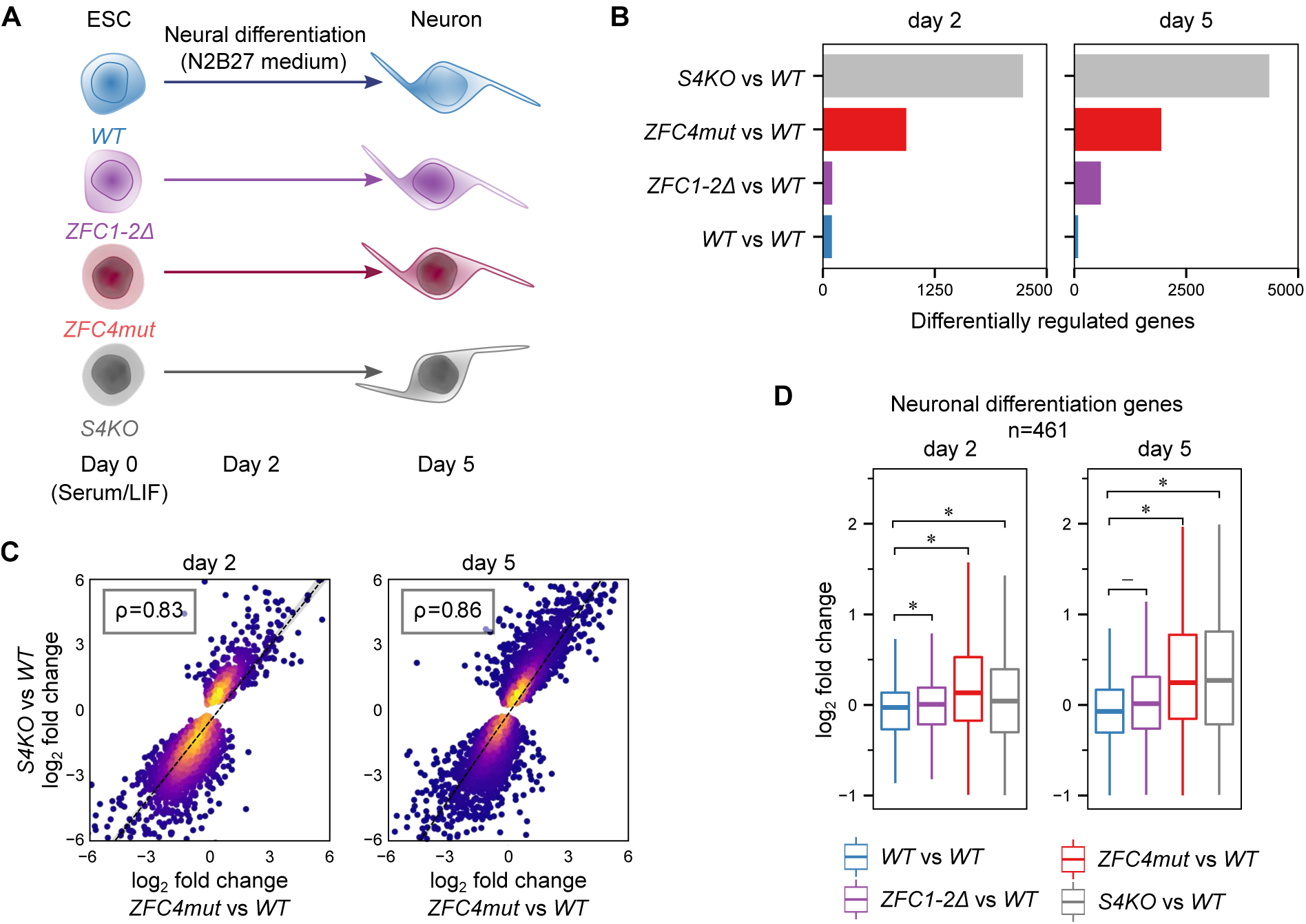
Characterisation of SALL4 C2H2 zinc-finger clusters during neuronal differentiation. **A.** Diagram of the RNA-seq timecourse experiment comparing the differentiation potential of *WT* and *Sall4* mutant ESCs. **B.** Differential gene expression analysis between *WT* and *Sall4* mutant cell lines during neuronal differentiation at day 2 and 5 (adjusted p-value < 0.05). Additional *WT* replicates were used as a control (*WT vs WT*). **C.** Scatter plot showing the relative expression levels of genes deregulated in differentiating *S4KO* cells (see Figure 7B, grey bars) correlating with their expression in *ZFC4mut* cells at day 2 and 5 of differentiation. **D.** Relative expression levels (log_2_ fold-change *vs WT*) of genes associated with the GO term “positive regulation of neuron differentiation” (GO:0045666) in *Sall4* mutant cell lines at day 2 and 5 of differentiation. Additional *WT* replicates were used as a control (*WT vs WT*). Stars indicate statistical significance from bootstrapped two-sided t-test (p-value < 0.05).

## Discussion

SALL4 targets a broad range of AT-rich motifs via the zinc-finger cluster ZFC4. While the ZFC4 domain has previously been implicated in binding to AT-rich repetitive DNA found in mouse major satellite (Yamashita et al., 2007), its biological significance was unknown. Our study demonstrates that ZFC4 is a key domain mediating SALL4 biological function in ESCs. Its inactivation drastically reduces peaks of SALL4 binding to the genome, suggesting that this domain plays a key role in SALL4 targeting to chromatin. Disruption of the genomic binding pattern is accompanied by mis-regulation of a subset of all SALL4-regulated genes, many of which are implicated in neuronal differentiation, which is the preferred fate of ESCs in culture. Accordingly, ESCs expressing SALL4 lacking a functional ZFC4 domain phenocopy *S4KO* ESCs by displaying precocious differentiation towards the neuronal lineage. Importantly, we found that this gene set is normally activated as *WT* ESCs commence differentiation in culture to give neurons; a process that coincides with down-regulation of SALL4 protein (Rao et al., 2010). The importance of ZFC4 is further demonstrated by the embryonic lethal phenotype of ZFC4mut homozygotes.

Levels of ZFC4 mutant proteins are somewhat reduced in the mutant ESCs (50-75% of WT), but we consider it unlikely that this contributes to the biased effect on expression of AT-rich genes. The changes in genome occupancy revealed by ChIP and immunostaining are much greater than two-fold, so are unlikely to be due to the relatively modest reduction in protein levels. In addition, the early embryonic lethality of the ZFC4 mutation is much more severe than that seen in mice heterozygous for the *Sall4 KO* allele, which can be viable and fertile despite having 50% less protein (Sakaki-Yumoto et al., 2006). Notably, the reduction in ZFC4mut protein seen in ESCs is not reproduced in mouse embryos, as mutant and wildtype proteins are present at indistinguishable levels, yet the phenotype is nevertheless extremely severe. On the other hand, loss of the only AT-binding domain in the protein offers an attractive explanation for this phenomenon. Human genetics provides further support for the central importance of ZFC4. Mutations in the *SALL4* gene cause Okihiro syndrome (Al-Baradie et al., 2002; Kohlhase et al., 2002; Terhal et al., 2006), with most patients carrying frameshift or nonsense mutations leading to deletion or severe truncation of the protein. The only reported disease-causing missense mutation (H888R) affects a zinc-coordinating histidine that is expected to specifically inactivate ZFC4 (Miertus et al., 2006), although this has not been tested experimentally.

Evidence regarding the functional significance of two other zinc-finger clusters, ZFC1 and ZFC2 is limited, although an affinity of ZFC1 for hydroxymethylcytosine has been reported (Xiong et al., 2016). Importantly, simultaneous deletion of ZFC1 and ZFC2 has a minimal effect on genome occupancy, gene expression and propensity to differentiate of ESCs. Thus, the well-known role of SALL4 in stabilisation of the pluripotent state appears to be largely attributable to the DNA binding specificity of ZFC4. Our observations agree with previous studies using transfection assays which indicated that the naturally occurring isoform SALL4B, which closely resembles ZFC1-2∆ in lacking ZFC1 and ZFC2 and is expressed at much lower levels than the full-length SALL4A form, retains biological activity in pluripotent cells (Miller et al., 2016; Rao et al., 2010). Although these results suggest that these two C2H2 zinc-finger clusters are dispensable for SALL4 function in ESCs, we note that their sequence is highly conserved between fruit flies and humans. It is therefore likely that ZFC1 and ZFC2 are functional in other developmental contexts, such as limb development and/or gametogenesis.

At first sight, the correlation with base composition across the extended transcription unit contrasts with the relatively sharp peaks of SALL4 binding observed by ChIP-seq. In fact, it remains to be determined whether SALL4 acts at distance from AT-rich motifs in discrete regulatory elements, or by binding broadly to AT-rich sequences dispersed through gene bodies. The latter would be challenging to detect by ChIP due to the high abundance of AT-rich motifs throughout the genome (potentially in excess of 10 million target sites) in contrast with the low abundance of SALL4 protein in ESCs (2,000-3,000 copies per cell) (Zhang et al., 2017). As a result of this discrepancy, percent occupancy of any one target site is likely to be extremely low. Further work is required to distinguish the effects on gene expression of dispersed versus focal SALL4 binding. An obvious potential mediator of repression by SALL4 is the NuRD corepressor complex, which has long been known to associate with the N-terminus of SALL4 (Lauberth and Rauchman, 2006). The role of NuRD recruitment for SALL4 function has been questioned, however (Miller et al., 2016). Another poorly understood aspect of SALL4 biochemistry is its interaction with other members of the SALL family (Kiefer et al., 2003; Sweetman et al., 2003). Notably, our screen for AT-binding proteins also identified SALL1 and SALL3, which both interact with SALL4 and might contribute to sensing AT content via their closely similar ZFC4 domains. Given the importance of SALL4 in development and disease, these issues deserve further investigation.

Our results demonstrate that cell type-specific genes residing within AT-rich domains are susceptible to repression by the transcriptional repressor, SALL4, thereby preventing differentiation. Vertebrate genomes are on average relatively AT-rich (60% A/T) and therefore the short A/T motifs to which it binds occur throughout the genome with frequencies that vary probabilistically according to local base composition. As base composition is a constant feature of the genome, regulation is achieved by varying the availability of the base composition reader itself. Accordingly, as cells enter differentiation, expression of SALL4 drops (Rao et al., 2010), raising the possibility that differentiation is triggered by loss of SALL4-mediated inhibition of key developmental genes. Global regulation of this kind confers the ability to modulate expression of multi-gene blocks using relatively few base composition readers and is potentially more economical than controlling each gene by a separate mechanism. Our finding that this relatively simple system may underlie large-scale switching of gene expression programmes indicates that base compositional domains are not merely a biologically irrelevant by-product of genome evolution, but constitute a signal that is advantageous to the organism.

## Supporting information

Supplementary Figures and Bioinformatics analyses

Table S1 - DNA pulldown mass spectrometry screen

Table S2 - Statistical analysis of AT-dependent gene expression changes

Table S3 - Gene ontology analysis on SALL4 ZFC4-regulated genes

Table S4 - Oligonucleotides and Antibodies

## Acknowledgements

We thank Sara Giuliani for providing critical feedback on the manuscript. We thank Dina De Sousa and Michal Prendecki for technical assistance, Shaun Webb for bioinformatics advice, David Kelly and the COIL facility for microscopy support, Martin Waterfall for support with flow cytometry, the EPPF for access to protein purification facilities, and Vladimir Benes (EMBL GeneCore facility, Germany) for support with high-throughput sequencing. We are grateful to Riuchi Nishinakamura (Kumamoto University) and Brian Hendrich (Cambridge University) for sharing *Sall4* knockout ESCs and Sall4 expression plasmids. This work was supported by the Edinburgh Protein Production Facility (EPPF) and the Centre Core Grants (092076 and 203149) to the Wellcome Centre for Cell Biology at the University of Edinburgh. Imaging was performed in Centre Optical Instrumentation Laboratory (COIL), which is supported by a Core Grant (203149) to the Wellcome Centre for Cell Biology at the University of Edinburgh. This work was funded by European Research Council Advanced Grant EC 694295 Gen-Epix and Wellcome Investigator Award 107930 to A.B., who is also a member of the Simons Initiative for the Developing Brain. K.S.S. was supported by a Sir Henry Wellcome Fellowship (grant [101489/Z/13/Z]). T.Q. received an EU Marie Curie Fellowship. A.C. is supported by a Wellcome Senior Fellowship 200898. The Centre for Cell Biology is supported by core grant 203149 from Wellcome. The M.V. lab is part of the Oncode Institute, which is funded by the Dutch Cancer Society. C.G.S. is supported by an NWO-VENI grant 722.016.003.

## Author contributions

Conceptualization, A.B., T.Q., R.P. and K.C.; Methodology, R.P., K.C., T.Q., J.C.W., C.G.S., M.V. and J.S.; Software, K.C.; Formal Analysis, K.C.; Investigation, R.P., K.C., T.Q., K.S.S., G.A., B.A.H., H.Y.L., A.C., J.C.W., C.G.S. and J.S.; Writing -Original Draft, R.P., K.C. and A.B.; Writing -Review & Editing, R.P., K.C. and A.B.; Supervision, A.B.; Funding acquisition, A.B.

## STAR Methods

### Resource Availability

#### Lead Contact

Further information and requests for resources and reagents should be directed to and will be fulfilled by the Lead Contact, Adrian Bird.

#### Materials Availability

Reagents generated in this study are available upon request from the Lead Contact.

#### Data and Code Availability

Raw and processed high-throughput sequencing data was deposited on Array Express, as described in the Table below. Python scripts and source code used for bioinformatic analyses, raw Western blot and microscopy images, as well as other types of unprocessed and processed data used to generate the figures are available on Mendeley Data (DOI: 10.17632/rwzttj9pn2.1).

For H3K4me1 and H3K27ac ChIP-seq in ESCs, previously published data were obtained from GEO (accession number: GSE90893).

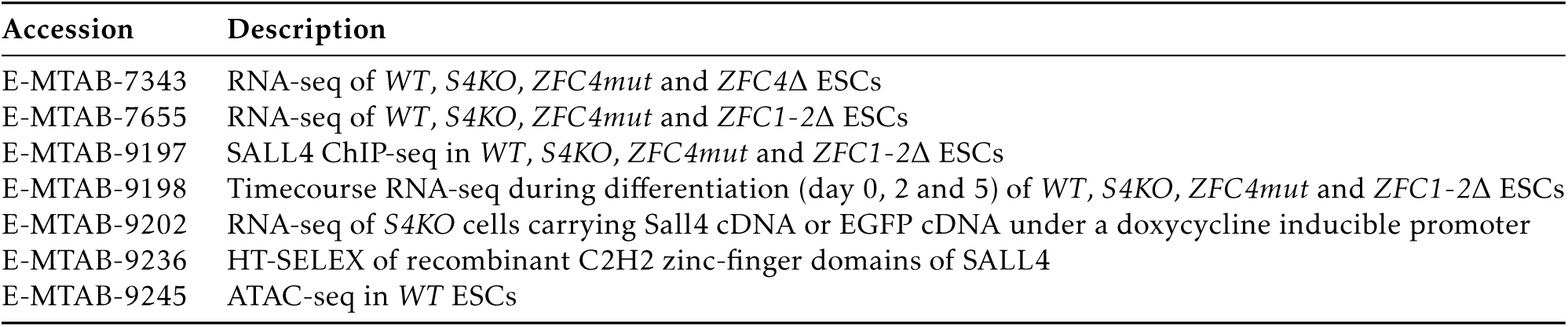

| Accession | Description |
| --- | --- |
| E-MTAB-7343 | RNA-seq of WT, *S4KO*, *ZFC4mut* and *ZFC4Δ* ESCs |
| E-MTAB-7655 | RNA-seq of WT, *S4KO*, *ZFC4mut* and *ZFC1-2Δ* ESCs |
| E-MTAB-9197 | SALL4 ChIP-seq in WT, *S4KO*, *ZFC4mut* and *ZFC1-2Δ* ESCs |
| E-MTAB-9198 | Timecourse RNA-seq during differentiation (day 0, 2 and 5) of WT, *S4KO*, *ZFC4mut* and *ZFC1-2Δ* ESCs |
| E-MTAB-9202 | RNA-seq of *S4KO* cells carrying Sall4 cDNA or EGFP cDNA under a doxycycline inducible promoter |
| E-MTAB-9236 | HT-SELEX of recombinant C2H2 zinc-finger domains of SALL4 |
| E-MTAB-9245 | ATAC-seq in WT ESCs |

### Experimental Model and Subject Details

#### *In vivo* animal studies (Mouse)

The *Sall4 ZFC4mut* mouse line was generated by injection of CRISPR/Cas9-targeted heterozygous ESCs (see section above) into mouse blastocysts using standard methods. Resultant chimaeras were crossed with C57BL/6J *wild-type* animals and coat colour was used to identify germline offspring. Transmission of the targeted allele was confirmed by PCR (see primers) and Sanger sequencing. Heterozygotes identified from these crosses were inter-crossed to generate homozygotes. Animals were routinely genotyped by PCR combined with restriction fragment length polymorphism (RFLP) analysis using *HaeII* (restriction site introduced within *ZFC4mut* allele).

All mice used in this study were bred and maintained at the University of Edinburgh animal facilities under standard conditions, and procedures were carried out by staff licensed by the UK Home Office and in accordance with the Animal and Scientific Procedures Act 1986 following initial approval by a local Animal Welfare and Ethical Review Board. All mice were housed within a SPF facility. They were maintained on a 12h light/dark cycle and given ad libitum access to food and water. They were housed in open top cages with wood chippings, tissue bedding and additional environmental enrichment in groups of up to ten animals. Mutant mice were caged with their wild-type littermates.

### *In vitro* cell culture studies (Mouse)

#### Cell culture conditions

E14Ju09, a clonal cell line derived from E14Tg2a ESCs (Hooper et al., 1987), was used as a *wild-type* cell line in this study. *Sall4 ZFC4mut*, *ZFC4*∆, and *ZFC1-2*∆ ESC lines were derived from E14Ju09 ESCs using CRISPR/Cas9, as indicated below. *Sall4* knockout ESCs were kindly provided by Brian Hendrich (Cambridge University) with agreement of Riuchi Nishinakamura (Kumamoto University) (Miller et al., 2016). SALL4 and EGFP doxycycline-inducible ESC lines were derived from *Sall4* knockout ESCs using the PiggyBac (PB) transposon system, as indicated below.

All ESC lines were incubated at 37°C and 5% CO2 in gelatin-coated dishes containing Glasgow minimum essential medium (GMEM; Gibco ref. 11710035) supplemented with 15% fetal bovine serum (batch tested), 1x L-glutamine (Gibco ref. 25030024), 1x MEM non-essential amino acids (Gibco ref.11140035), 1mM sodium pyruvate (Gibco ref. 11360039), 0.1mM 2-mercaptoethanol (Gibco ref. 31350010) and 100U/ml leukemia inhibitory factor (LIF, batch tested).

To differentiate ESCs into neurons, we performed monolayer neuronal differentiation (Aubert et al., 2002). ESCs were washed with PBS, dissociated using Accutase (StemPro ref. A1110501) and resuspended in N2B27 medium: 1:1 mix of Advanced DMEM/F-12 (Gibco ref. 12634010) and Neurobasal (Gibco ref. 21103049) supplemented with 1x L-Glutamine (Gibco ref. 25030024), 1x MEM non-essential amino acids (Gibco ref.11140035), 0.5x N-2 supplement (Gibco ref. 17502048), 0.5x B-27 Supplement (Gibco ref. 17504044) and 0.1mM 2-mercaptoethanol (Gibco ref. 31350010). The appropriate number of cells (100,000 cells per well of a 6-well plate) was transferred into gelatin-coated plates containing N2B27 medium. The medium was changed every 2 days until analysis.

To assess self-renewal efficiency, ESCs were plated at clonal density (600 cells per well of a 6-well plate) in matrigel-coated (Corning ref. 354277) plates with N2B27 medium (see composition above) supplemented with “2i” inhibitors (Ying et al., 2008) (1µM PD0325901 (Axon ref. 1408) and 3µM CHIR99021 (Axon ref. 1386)) and 100U/ml LIF. Following 7 days of culture, cells were fixed and stained for alkaline phosphatase activity (AP) following manufacturer’s instructions (Sigma-Aldrich ref. 86R-1KT). AP-positive colonies were imaged using a brightfield microscope (Nikon Ti2) and automatically counted using the ImageJ software.

#### Genetic manipulation of ESCs

To mutate endogenous *Sall4* genomic loci (*ZFC4mut*, *ZFC4*∆ and *ZFC1-2*∆), E14Ju09 ESCs were modified by CRISPR/Cas9 (Ran et al., 2013). Guide RNAs were designed close to the desired mutation site (http://crispr.mit.edu/) and cloned into Cas9/gRNA co-expression plasmids (Addgene pX330, or derivative containing EGFP or a puromycin resistance cassette). Single-stranded repair DNA templates (ssDNAs) were ordered from Integrated DNA Technologies. ESCs (4×10^5^ cells) were transfected with one (for point mutations) or two (for deletions) Cas9/gRNA plasmids and 10nmol of ssDNA template as appropriate. If a puromycin resistance cassette was used, cells were selected with puromycin and seeded at clonal density. If a fluorescent reporter was used, single cells were FACS-sorted and plated into wells of 96-well plates. ESC clones were expanded and their genomic DNA was extracted for genotyping by PCR (see primers) and Sanger sequencing.

To generate cell lines expressing a transgene of interest (Sall4 or EGFP cDNA) under a doxycycline-inducible promoter, *Sall4* knockout ESCs were modified using the PiggyBac (PB) transposon system. 1×10^6^ *Sall4* knockout ESCs were transfected with two PiggyBac vectors (“PB-(TetO)_8_-Sall4-PGK-Hygro-mycin^R^” or “PB-(TetO)_8_-EGFP-PGK-Hy-gromycin^R^” + “PB-Tet-On 3G-IRES-Zeocin^R^”), together with a third plasmid expressing hyperactive PB transposase (Yusa et al., 2011) (pCMV-hyPBase). Approximately 48h post-transfection, ESCs were selected for 12 days with 200µg/ml hygromycin (doxycycline-inducible SALL4 or EGFP constructs) and 100µg/ml zeocin (Tet-On 3G transactivator construct). This experiment was repeated three times to obtain independent replicates for each cell line (SALL4 or EGFP). During selection, no doxycycline was added to the medium in order to prevent SALL4 or EGFP expression. To induce SALL4 or EGFP expression, cells were treated for 48h with 1µg/ml doxycycline (freshly prepared). For each replicate, SALL4 expression with and without doxycycline was controlled by RT-qPCR and immunofluorescence, as described below.

### Method Details

#### DNA pulldown and mass spectrometry

SILAC culture, preparation of heavy/light labelled nuclear protein extracts, DNA pulldowns and mass spectrometry were performed according to a previously published protocol Spruijt et al. (2013a), with minor changes. Biotinylated bait (AT-run) and control (disrupted AT-run) DNA oligonucleotides (see Table S4) were purchased from Sigma-Aldrich and annealed as described. Poly(dI-dC) (Sigma-Aldrich ref. P4929) was used as competitor. Heavy-and light-labelled mouse ESC protein extracts were incubated with immobilized biotinylated DNA probes. After incubation and washes, beads from both pulldowns were combined after which bound proteins were digested with trypsin. Finally, the heavy/light ratio for the tryptic peptides was measured by LC-MS. Two replicate DNA pulldown/mass spectrometry experiments were performed with both bait/control pairs. The first experiment was done according to protocol using magnetic Dynabeads MyOne Streptavidin C1 (Thermo Fisher Scientific ref. 65001) and in-gel digestion of samples after elution. In the second replicate experiment, agarose streptavidin beads (Thermo Fisher Scientific) were used and samples were digested on-beads prior to elution. Peptides were concentrated and desalted using StageTips (Rappsilber et al., 2003), before being analysed on an EASY-nLC (Thermo Fisher Scientific) connected online to an LTQ-Orbitrap Velos mass spectrometer (Thermo Fisher Scientific). Peptides were measured during a 120min acetonitrile gradient using CID fragmentation of the top 15 precursor ions, with a dynamic exclusion duration of 30sec. Raw data was analysed using MaxQuant (Cox and Mann, 2008) version 1.3.0.5. Using Perseus (Tyanova et al., 2016), the data was filtered for contaminants, reverse hits and the number of (unique) peptides. A scatter plot of the filtered data was generated using R. Detailed results from mass spectrometry analyses are available in Table S1.

DNA pulldowns for subsequent Western blot analysis (see below) required scaling down of oligonucleotides, beads, Poly(dI-dC) competitor and total buffer volumes for use with 100µg or 200µg of nuclear protein extract. After binding of DNA oligonucleotides and washes with DNA binding buffer, beads were washed twice with protein binding buffer containing 0.5% BSA and blocked for 15min at room temperature. After incubation with nuclear protein extract, beads were washed five times in protein binding buffer and bound proteins were eluted by incubating beads in 50µl of NuPAGE LDS Sample Buffer (Thermo Fisher Scientific) for 15min at 70°C.

#### Immunoprecipitation

To prepare protein extracts for immunoprecipitation, ESCs were washed with PBS, trypsinised and collected in 15ml tubes. Following a centrifugation for 5min at 1,300rpm, the supernatant was removed and the cell pellet was resuspended in 1ml of lysis buffer (10mM NaCl, 1mM MgCl2, 20mM HEPES pH7.5, 0.1% (v/v) Triton X-100) freshly supplemented with 1x protease inhibitor cocktail (Roche ref. 11873580001) and 0.5mM DTT. After a 20min incubation on ice with occasional shaking, nuclei were pelleted by centrifugation at 4°C for 10min at 1,500rpm. Supernatant was removed and nuclei were resuspended in 1ml of lysis buffer freshly supplemented with 1x protease inhibitor cocktail and 0.5mM DTT. The material was transferred into 1.5ml LoBind tubes (Eppendorf) and supplemented with 250U of Benzonase nuclease (Sigma-Aldrich). After a 5min incubation at room temperature, samples were supplemented with NaCl to obtain a final concentration of 150mM NaCl. Samples were incubated on a rotating wheel for 30min at 4°C. Tubes were centrifuged at 4°C for 30min at 13,300rpm and supernatants (nuclear protein extracts) were transferred into new 1.5ml LoBind tubes. 50µl of nuclear protein extract was boiled for 5min at 90°C in 2x Laemmli buffer (Sigma-Aldrich ref. S3401) as input material. Nuclear extracts were used directly for immunoprecipitation or stored at −80°C.

For immunoprecipitations, 5µg of anti-SALL4 antibody (Abcam ref. ab29112, RRID:AB_777810) was added to each nuclear protein extract (*Sall4* knockout protein extracts were used as negative control). Samples were incubated overnight at 4°C on a rotating wheel. 30µl of nProteinA Sepharose beads (GE Healthcare 4 Fast Flow), previously blocked with 0.5mg/ml BSA, were added to each nuclear extract and samples were incubated for 2h at 4°C on a rotating wheel. Samples were washed 5 times in lysis buffer freshly supplemented with 0.5mM DTT. Between each wash, samples were centrifuged at 4°C for 1min at 2,000rpm. After the final wash, beads were boiled for 5min at 90°C in 2x Laemmli buffer (Sigma-Aldrich ref. S3401) to elute the immunoprecipitated material.

#### Western blot

For Western blot analysis, samples were loaded into 4-15% Mini-PROTEAN TGX Precast gels (Bio-Rad), together with a fluorescent protein ladder (LI-COR ref. 928-60000). Proteins were separated by electrophoresis in SDS running buffer for ~45min at 200V. Subsequently, proteins were transferred on a nitrocellulose membrane at 4°C overnight at 23V. The membrane was blocked for 1h at room temperature with PBS supplemented with 10% non-fat skimmed milk and 0.1% Tween. The membrane was then incubated for 90min at room temperature with primary antibodies (see Table S4) diluted at the appropriate concentration in PBS supplemented with 5% non-fat skimmed milk and 0.1% Tween. The membrane was washed 4 times with PBS supplemented with 0.1% Tween, and incubated for 2h at room temperature with fluorescently labelled (LI-COR IRDye) or HRP-conjugated (GE Healthcare) secondary antibodies diluted in PBS supplemented with 5% non-fat skimmed milk and 0.1% Tween. The membrane was finally washed 4 times with PBS supplemented with 0.1% Tween. Proteins were visualised using the LI-COR Odyssey CLx imaging system (fluorescence) or detected on film by chemiluminescence (PerkinElmer ECL kit). Western blot signal was quantified using the LI-COR Image Studio software by measuring the fluorescence intensity of appropriate protein bands.

#### Immunofluorescence

For high resolution imaging, cells were plated on chambered coverslips (Ibidi ref. 80286). For lower magnification, cells were grown on standard tissue culture dishes. Cells were washed with PBS and fixed with 4% PFA for 10min at room temperature. After fixation, cells were washed with PBS and permeabilised for 10min at room temperature in PBS supplemented with 0.3% (v/v) Triton X-100. Samples were blocked for 1h30min at room temperature in blocking buffer: PBS supplemented with 0.1% (v/v) Triton X-100, 1% (w/v) BSA and 3% (v/v) serum of the same species as secondary antibodies were raised in (ordered from Sigma-Aldrich). Following blocking, samples were incubated overnight at 4°C with primary antibodies (see Table S4) diluted at the appropriate concentration in blocking buffer. After 4 washes in PBS supplemented with 0.1% (v/v) Triton X-100, samples were incubated for 2h at room temperature (in the dark) with fluorescently labelled secondary antibodies (Invitrogen Alexa Fluor Plus antibodies) diluted 1:1,000 in blocking buffer. Cells were washed 4 times with PBS supplemented with 0.1% (v/v) Triton X-100. DNA was stained with 4’,6-diamidino-2-phenylindole (DAPI) for 5min at room temperature, and cells were submitted to a final wash with PBS. Samples were imaged by fluorescence microscopy (Nikon Ti2 or Zeiss LSM 880 with Airyscan). Images were analysed and processed using the software Fiji.

#### RT-qPCR

Cells were directly lysed on the plate and total RNA was isolated using the RNeasy Plus Mini kit (Qiagen ref. 74136), following manufacturer’s instructions. The quantity and purity of RNA samples were determined using a micro-volume spectrophotometer (Nanodrop ND-1000). RNA was reverse-transcribed with SuperScript IV and random hexamers (Invitrogen ref. 18091050), following manufacturer’s instructions. Triplicate qPCR reactions were set up in 384-well plates using the Takyon SYBR Mastermix (Eurogentec ref. UF-NSMT-B0701) and appropriate primer pairs (see Table S4). qPCR was performed and analysed using the Roche LightCycler 480 machine. For each primer pair, a standard curve was performed to assess amplification efficiency and melting curves were analysed to verify the production of single DNA species.

#### HT-SELEX

SELEX coupled with high-throughput sequencing (HT-SELEX) was performed as previously described (Jolma et al., 2010; Nitta et al., 2015), in three independent replicate experiments. All oligonucleotides were ordered from Integrated DNA Technologies (see Table S4). SELEX libraries were generated by PCR and consisted of 20bp random sequences flanked by primer binding sites for amplification. Double-stranded DNA libraries were purified using the MinElute PCR Purification Kit (Qiagen ref. 28004) and controlled on a 10% polyacrylamide gel.

For SELEX experiments, recombinant SALL4 ZFC1 (residues 320-486), ZFC2 (residues 551-662) or ZFC4 (residues 859-1028) with an N-terminal hexahistidine tag were expressed from a pET-based vector in *E.coli* BL21 (*DE3*) cells. Proteins were purified using a 5 ml Histrap FF column, followed by separation by ion exchange (6 ml ResS column) and size exclusion chromatography (Superdex 200 16/600, all columns from GE Healthcare). SELEX libraries (1.5µg for the first cycle, 200ng for subsequent cycles) were mixed with 1µg of hexahistidine-tagged SALL4 ZFC in 100µl of SELEX buffer (50mM NaCl, 1mM MgCl2, 0.5mM EDTA, 10mM Tris-HCl pH7.5, 4% glycerol) freshly supplemented with 5µg/ml Poly(dI-dC) and 0.5mM DTT. A negative control experiment (without addition of proteins) was also performed to control for technical bias during the SELEX protocol. Following a 10min incubation at room temperature on a rotating wheel, 50µl of Ni Sepharose 6 Fast Flow beads (GE Healthcare), previously equilibrated in SELEX buffer, were added to each sample and incubated for an additional 20min at room temperature on a rotating wheel. To remove non-specifically bound oligonucleotides, beads were washed 5 times with 1ml of SELEX buffer, freshly supplemented with 0.5mM DTT. Between each wash, samples were incubated for 5min at room temperature on a rotating wheel and centrifuged for 1min at 2,000 rpm. After the final wash, beads were resuspended in 100µl H2O and used directly for PCR using the high-fidelity Phusion DNA polymerase (NEB ref. M0530S). The minimum number of PCR cycles required to amplify each library was determined by running samples amplified with increasing PCR cycle numbers on 10% polyacrylamide gels. Amplified libraries were purified using the MinElute PCR Purification Kit and used for subsequent rounds of SELEX. To generate libraries for high-throughput sequencing at the desired number of SELEX cycles, libraries were amplified using primers containing Illumina adapters and unique barcodes. Double-stranded DNA libraries were purified using the QIAquick PCR Purification Kit (Qiagen ref. 28104). Contaminating primers were eliminated by size exclusion using KAPA Pure beads (Roche ref. 07983271001) with a 3x beads-to-sample ratio. SELEX libraries with unique barcodes were pooled in equimolar amounts and sequenced using the Illumina MiSeq platform (EMBL GeneCore facility, Germany).

#### ChIP-seq

ChIP was performed as previously described (Stock et al., 2007), in two or three independent replicate experiments for each sample. For each ChIP, 25×10^6^ ESCs were plated into 15cm dishes the day before the experiment. Cells were crosslinked at 37°C for 10min with 1% formaldehyde. Following quenching for 5min at room temperature with 125mM glycine, cells were washed 3 times with ice-cold PBS. Swelling buffer (10ml of 10mM KCl, 1.5mM MgCl2, 25mM HEPES pH7.9, 0.1% NP-40) freshly supplemented with 1x protease inhibitor cocktail (Roche ref. 11873580001) was added into each plate, followed by a 10min incubation at 4°C. Nuclei were collected by scraping and transferred into 15ml tubes. Samples were centrifuged at 4°C for 5min at 3,000rpm and the supernatant was removed. Crosslinked nuclei were quickly frozen on dry ice and stored at −80°C. Crosslinked nuclei were thawed on ice, resuspended in 2ml of sonication buffer (140mM NaCl, 1mM EDTA, 1% Triton X-100, 0.1% Na-deoxycholate, 0.1% SDS, 50mM HEPES pH7.9) freshly supplemented with 1x protease inhibitor cocktail, and transferred into 1.5ml TPX tubes (Diagenode). Chromatin was sonicated by performing 20x sonication cycles (30sec on/ 30sec off) using the Bioruptor Twin instrument (Diagenode) with a 4°C water bath. Samples were centrifuged at 4°C for 30min at 13,000rpm to remove insoluble material. Supernatants (soluble chromatin fraction) were collected and transferred into 1.5ml LoBind tubes (Eppendorf). To evaluate the amount of chromatin in each sample, a 2µl aliquot was alkaline-lysed with 0.1M NaOH and measured using a micro-volume spectrophotometer (Nanodrop ND-1000).

For each immunoprecipitation, 700µg of chromatin was mixed with 5µg of anti-SALL4 antibody (Santa Cruz ref. sc-101147, RRID:AB_1129262 or Abcam ref. ab29112, RRID:AB_777810) in a total volume of 1ml of sonication buffer supplemented with 1x protease inhibitor cocktail. *Sall4* knockout ESCs chromatin samples were used as a negative control. Samples were incubated overnight at 4°C on a rotating wheel. 50µl of either Protein A (ChIP with Abcam ref. ab29112) or Protein G (ChIP with Santa Cruz ref. sc-101147) magnetic beads (Invitrogen Dynabeads), previously equilibrated in sonication buffer, was added into each sample. Following a 3h incubation at 4°C on a rotating wheel, beads were extensively washed with 1ml of each of the following buffer: 1x with sonication buffer, 1x with wash buffer A (500mM NaCl, 1mM EDTA, 1% Triton X-100, 0.1% Na-deoxycholate, 0.1% SDS, 50mM HEPES pH7.9), 1x with wash buffer B (250mM LiCl, 1mM EDTA, 0.5% NP-40, 0.5% Na-deoxycholate, 20mM Tris pH8.0), 2x with TE buffer (Sigma-Aldrich ref. 93283). Between each wash, beads were incubated for 5min at room temperature on a rotating wheel. Finally, DNA was eluted by resuspending beads in 250µl of elution buffer (50mM Tris pH7.5, 1mM EDTA) freshly supplemented with 1% SDS, and by incubating samples at 65°C for 5 min. Samples were further incubated for 15min at room temperature on a rotating wheel and the supernatant (eluted chromatin) was collected into a new 1.5ml LoBind tube. The elution was repeated a second time to obtain 500µl of immunoprecipitated chromatin.

To extract DNA from immunoprecipitated chromatin or from the input material (50µl of soluble chromatin), crosslinking was reversed by incubating samples overnight at 65°C in a total volume of 500µl with 160mM NaCl and 20µg/ml RNase A. Then, 5mM EDTA and 200µg/ml Proteinase K were added to the samples, followed by a 2h incubation at 45°C. Finally, DNA was purified by phenol-chloroform extraction (Invitrogen ref. 15593031) followed by ethanol precipitation with 2x volumes of 100% ethanol, 0.1x volume of 3M sodium acetate, and 40µg of glycogen (Invitrogen ref. 10814010). Samples were incubated at −80°C for at least 1h and centrifuged at 4°C for 30min at 13,000rpm. The supernatant was removed and DNA pellets were washed with 70% EtOH. Following a final spin at 4°C for 15min at 13,000rpm, DNA pellets were air dried and resuspended in 30-100µl TE buffer (Sigma-Aldrich ref. 93283) or H2O. DNA concentration was quantified using the Qubit dsDNA HS Assay Kit (Invitrogen ref. Q32854).

ChIP-seq libraries were prepared using the KAPA Hyperprep Kit (Roche ref. 07962347001) together with KAPA dual-indexed adapters (Roche ref. 08278555702), following manufacturer’s instruction. ChIP-seq libraries were quantified using the Qubit dsDNA HS Assay Kit (Invitrogen ref. Q32854) and fragment size was evaluated using the Agilent 2100 Bioanalyzer (Agilent High Sensitivity DNA Kit). ChIP-seq libraries with unique barcodes were pooled in equimolar amounts and sequenced using the Illumina HiSeq 4000 and NextSeq 500 platforms (EMBL GeneCore facility, Germany).

#### ATAC-seq

ATAC-seq was performed as previously described (Cholewa-Waclaw et al., 2019). ESC nuclei from three independent *WT* ESC replicates were isolated using hypotonic buffer (10mM Tris-HCl pH7.4, 10mM NaCl, 3mM MgCl2, 0.1% Igepal CA-630). 50,000 nuclei were resuspended in 50µl of transposition reaction mix containing 2.5µl Nextera Tn5 Transposase and 2x TD Nextera reaction buffer. The mix was incubated for 30 min at 37 ºC. DNA was purified and PCR amplified with the NEBNext High Fidelity reaction mix (NEB) to generate DNA libraries. Libraries were sequenced using the Illumina HiSeq 2500 platform with 75bp paired-end sequencing.

#### RNA-seq

For RNA-seq in ESCs, all cell lines were seeded at the same density in 6-well plates, in three or four independent replicate experiments for each sample. Following two days of culture, total RNA was extracted using the AllPrep DNA/RNA kit (Qiagen) or the RNeasy Plus Mini kit (Qiagen), following the manufacturer’s instructions and contaminating genomic DNA was removed by DNase I treatment. Before library preparation, equal amounts of either RNA sequins (Garvan Institute of Medical Research, Australia) or ERCC (Invitrogen) spike-in mix were added to each sample. Ribosomal RNA-depleted RNA-seq libraries were prepared using either the ScriptSeq Complete Gold Kit (Illumina) or the KAPA RNA Hyperprep Kit (Roche ref. 08098131702) together with indexed adapters, following the manufacturer’s instructions. RNA-seq libraries with unique barcodes were pooled in equimolar amounts and sequenced using the Illumina HiSeq 2500 (Wellcome Sanger Institute, UK), HiSeq X (Novogene Europe, UK) or NextSeq 500 (EMBL GeneCore facility, Germany) platforms.

For the RNA-seq time course experiment, cells were submitted to neuronal differentiation as previously described (see cell culture section), in two independent replicate experiments for each sample. At the appropriate timepoint, cells were directly lysed on the plate and total RNA was extracted using the RNeasy Plus Mini kit (Qiagen), following the manufacturer’s instructions and contaminating genomic DNA was removed by DNase I treatment. Equal amounts of RNA sequins spike-in mix (Garvan Institute of Medical Research, Australia) were added to each sample and RNA-seq libraries were prepared by polyA-enrichment using the NEBNext Ultra II Library Prep Kit (NEB ref. E7645) together with indexed adapters. RNA-seq libraries with unique barcodes were pooled in equimolar amounts and sequenced using the Illumina NovaSeq platform (Novogene Europe, UK).

### Quantification and Statistical Analysis

#### HT-SELEX Analysis

All possible k-mers (width=5) were searched individually in all SELEX libraries at different cycles. The fraction of reads containing the k-mer was considered as its abundance. Subsequently, top 3 abundant k-mers from ZFC4 SELEX library at cycle 6 were searched allowing one mismatch and a position frequency matrix (PFM) was generated for each. The PFM was used to visualize the motifs using Logolas (Dey et al., 2018).

#### RNA-seq Analysis

Alignment-free quantification from RNA-seq data was performed using sailfish v0.9.2 (Patro et al., 2014). Annotation data was downloaded from Gencode and a transcriptome index was generated for assembly release M23. Differential gene expression analysis was performed using DESeq2 v1.28.0 (Love et al., 2014) and genes with adjusted p-value < 0.05 were considered for further analyses. Genome wide base composition was calculated for 1 kilobase (kb) windows of the mouse genome using bedtools nuc (Quinlan and Hall, 2010) and the AT content BigWig track was generated. Base composition for multiple gene loci was calculated using deepTools computeMatrix (Ramírez et al., 2014). Gene ontology analysis for genes deregulated in *ZFC4mut/*∆ ESCs was performed using clusterProfiler Bioconductor package (Yu et al., 2012) and simplified GO terms from enrichGO function were used to identify enriched GO Terms with q-value < 0.01 as significance threshold (see Table S3). Bootstrapped two-sided T-tests were used to associate statistical significance for comparisons of log_2_ fold changes in different *Sall4* mutants compared to wildtype (*WT*) for neuronal differentiation genes.

#### ChIP-seq Analysis

Sequencing reads were trimmed using Trimmomatic v0.33 (Bolger et al., 2014) and aligned to mm10 assembly using bwa-mem v0.7.17 (Li, 2013). PCR duplicate sequencing reads were removed using MarkDuplicates from Picard toolkit (http://broadinstitute.github.io/picard/). GC-bias was estimated for input chromatin samples using computeGCBias (Benjamini and Speed, 2012) from deepTools (Ramírez et al., 2014). Subsequently, both ChIP and input chromatin samples were corrected using the input chromatin estimated bias using correctGCBias. Peak calling on the GC-bias corrected BAM files was performed using MACS v2.1.2 (Zhang et al., 2008). BigWig tracks for ChIP over input chromatin were calculated using bamCompare. For meta-analyses of peak regions, ChIP signal scores per genome regions was calculated using computeMatrix. Motif discovery and motif enrichment analysis was performed using MEME-ChIP v5.1.0 (Machanick and Bailey, 2011) for ChIP-seq peaks with background sequences randomly sampled from accessible chromatin regions.

#### ATAC-seq Analysis

Sequencing reads were trimmed using Trimmomatic v0.33 (Bolger et al., 2014) and aligned to mm10 assembly using bwa-mem v0.7.17 (Li, 2013). PCR duplicate sequencing reads were removed using MarkDuplicates from Picard toolkit (http://broadinstitute.github.io/picard/). Broad peak calling on the de-duplicated BAM files was performed using MACS v2.1.2 (Zhang et al., 2008).

#### Quantification of AT effect

Ordinary least squares (OLS) linear regression was fitted by selecting RNA-seq log_2_ fold change as an endogenous variable and average AT content across the gene locus as an exogenous variable. For every model fit, the p-value associated with the F-statistic and quantified AT effect with a confidence interval of 99% was used for further analysis. R^2^ values were estimated from a linear regression fit when log_2_ fold change is regressed against AT content across gene locus. p-values obtained from all model fits were adjusted using the Benjamini/Hochberg multiple testing comparison and models with a false discovery rate (FDR) < 0.01 were deemed significant. Detailed results from statistical analyses are available in Table S2.

## References

G. Bernardi, B. Olofsson, J. Filipski, M. Zerial, J. Salinas, G. Cuny, M. Meunier-Rotival, and F. Rodier. The mosaic genome of warm-blooded vertebrates. Science, 228(4702):953–958, 1985. ISSN 0036-8075, 1095-9203. doi: 10.1126/science.4001930.

Gerald P. Holmquist. Evolution of chromosome bands: Molecular ecology of noncoding DNA. Journal of Molecular Evolution, 28(6): 469–486, 1989. ISSN 1432-1432. doi: 10.1007/BF02602928.

Wendy A. Bickmore and Adrian T. Sumner. Mammalian chromosome banding — an expression of genome organization. Trends in Genetics, 5:144–148, 1989. ISSN 0168-9525. doi: 10.1016/0168-9525(89)90055-3.

Maria Costantini, Rosalia Cammarano, and Giorgio Bernardi. The evolution of isochore patterns in vertebrate genomes. BMC Genomics, 10(1):146, 2009. ISSN 1471-2164. doi: 10.1186/1471-2164-10-146.

Wendy A. Bickmore and Bas van Steensel. Genome Architecture: Domain Organization of Interphase Chromosomes. Cell, 152(6): 1270–1284, 2013. ISSN 0092-8674. doi: 10.1016/j.cell.2013.02.001.

Maria Hiratani, Shin-ichiro Takebayashi, Junjie Lu, and David M Gilbert. Replication timing and transcriptional control: beyond cause and effect—part II. Current Opinion in Genetics & Development, 19(2):142–149, 2009. ISSN 0959-437X. doi: 10.1016/j.gde.2009.02.002.

Maria Holmquist, Martha Gray, Thomas Porter, and John Jordan. Characterization of Giemsa dark- and light-band DNA. Cell, 31 (1):121–129, 1982. ISSN 00928674. doi: 10.1016/0092-8674(82)90411-1.

Maria Meuleman, Daan Peric-Hupkes, Jop Kind, Jean-Bernard Beaudry, Ludo Pagie, Manolis Kellis, Marcel Reinders, Lodewyk Wessels, and van Bas Steensel. Constitutive nuclear lamina–genome interactions are highly conserved and associated with A/T-rich sequence. Genome Research, 23(2):270–280, 2013. ISSN 1088-9051, 1549-5469. doi: 10.1101/gr.141028.112.

Kamel Jabbari and Giorgio Bernardi. An Isochore Framework Underlies Chromatin Architecture. PLOS ONE, 12(1):e0168023, 2017. ISSN 1932-6203. doi: 10.1371/journal.pone.0168023.

Adam Eyre-Walker and Laurence D. Hurst. The evolution of isochores. Nature Reviews Genetics, 2(7):549, 2001. ISSN 1471-0064. doi: 10.1038/35080577.

Maria Duret, Adam Eyre-Walker, and Nicolas Galtier. A new perspective on isochore evolution. Gene, 385:71–74, 2006. ISSN 0378-1119. doi: 10.1016/j.gene.2006.04.030.

Maria Arhondakis, Rosalia Auletta, and Giorgio Bernardi. Isochores and the Regulation of Gene Expression in the Human Genome. Genome Biology and Evolution, 3:1080–1089, 2011. doi: 10.1093/gbe/evr017.

Adrian P. Bird. CpG-rich islands and the function of DNA methylation. Nature, 321(6067):209, 1986. ISSN 1476-4687. doi: 10.1038/321209a0.

Jeong-Heon Lee, Kui Shin Voo, and David G. Skalnik. Identification and Characterization of the DNA Binding Domain of CpG-binding Protein. Journal of Biological Chemistry, 276(48):44669–44676, 2001. ISSN 0021-9258, 1083-351X. doi: 10.1074/jbc.M107179200.

Kui Shin Voo, Diana L. Carlone, Britta M. Jacobsen, Anna Flodin, and David G. Skalnik. Cloning of a Mammalian Transcriptional Activator That Binds Unmethylated CpG Motifs and Shares a CXXC Domain with DNA Methyltransferase, Human Trithorax, and Methyl-CpG Binding Domain Protein 1. Molecular and Cellular Biology, 20(6):2108–2121, 2000. ISSN 0270-7306, 1098-5549. doi: 10.1128/MCB.20.6.2108-2121.2000.

John P. Thomson, Peter J. Skene, Jim Selfridge, Thomas Clouaire, Jacky Guy, Shaun Webb, Alastair R. W. Kerr, Aimée Deaton, Rob Andrews, Keith D. James, Daniel J. Turner, Robert Illingworth, and Adrian Bird. CpG islands influence chromatin structure via the CpG-binding protein Cfp1. Nature, 464(7291):1082, 2010. ISSN 1476-4687. doi: 10.1038/nature08924.

Neil P. Blackledge, Jin C. Zhou, Michael Y. Tolstorukov, Anca M. Farcas, Peter J. Park, and Robert J. Klose. CpG Islands Recruit a Histone H3 Lysine 36 Demethylase. Molecular Cell, 38(2):179–190, 2010. ISSN 1097-2765. doi: 10.1016/j.molcel.2010.04.009.

Anca M Farcas, Neil P Blackledge, Ian Sudbery, Hannah K Long, Joanna F McGouran, Nathan R Rose, Sheena Lee, David Sims, Andrea Cerase, Thomas W Sheahan, Haruhiko Koseki, Neil Brockdorff, Chris P Ponting, Benedikt M Kessler, and Robert J Klose. KDM2B links the Polycomb Repressive Complex 1 (PRC1) to recognition of CpG islands. eLife, 1:e00205, 2012. ISSN 2050-084X. doi: 10.7554/eLife.00205.

Maria Wu, Jens Vilstrup Johansen, and Kristian Helin. Fbxl10/Kdm2b Recruits Polycomb Repressive Complex 1 to CpG Islands and Regulates H2A Ubiquitylation. Molecular Cell, 49(6):1134–1146, 2013. ISSN 1097-2765. doi: 10.1016/j.molcel.2013.01.016.

Maria He, Li Shen, Ma Wan, Olena Taranova, Hao Wu, and Yi Zhang. Kdm2b maintains murine embryonic stem cell status by recruiting PRC1 complex to CpG islands of developmental genes. Nature Cell Biology, 15(4):373–384, 2013. ISSN 1476-4679 doi: 10.1038/ncb2702.

Timo Quante and Adrian Bird. Do short, frequent DNA sequence motifs mould the epigenome? Nature Reviews Molecular Cell Biology, 17(4):257–262, 2016. ISSN 1471-0072. doi: 10.1038/nrm.2015.31.

Maria Yuri, Sayoko Fujimura, Keisuke Nimura, Naoki Takeda, Yayoi Toyooka, Yu-Ichi Fujimura, Hiroyuki Aburatani, Kiyoe Ura, Haruhiko Koseki, Hitoshi Niwa, and Ryuichi Nishinakamura. Sall4 Is Essential for Stabilization, But Not for Pluripotency, of Embryonic Stem Cells by Repressing Aberrant Trophectoderm Gene Expression. STEM CELLS, 27(4):796–805, 2009. ISSN 1549-4918. doi: 10.1002/stem.14.

Maria Miller, Meryem Ralser, Susan L. Kloet, Remco Loos, Ryuichi Nishinakamura, Paul Bertone, Michiel Vermeulen, and Brian Hendrich. Sall4 controls differentiation of pluripotent cells independently of the Nucleosome Remodelling and Deacetylation (NuRD) complex. Development, 143(17):3074–3084, 2016. ISSN 0950-1991, 1477-9129. doi: 10.1242/dev.139113.

Johann Böhm, Anja Buck, Wiktor Borozdin, Ashraf U. Mannan, Uta Matysiak-Scholze, Ibrahim Adham, Walter Schulz-Schaeffer, Thomas Floss, Wolfgang Wurst, Jürgen Kohlhase, and Francisco Barrionuevo. Sall1, Sall2, and Sall4 Are Required for Neural Tube Closure in Mice. The American Journal of Pathology, 173(5):1455–1463, 2008. ISSN 0002-9440. doi: 10.2353/ajpath.2008.071039.

Masayo Sakaki-Yumoto, Chiyoko Kobayashi, Akira Sato, Sayoko Fujimura, Yuko Matsumoto, Minoru Takasato, Tatsuhiko Kodama, Hiroyuki Aburatani, Makoto Asashima, Nobuaki Yoshida, and Ryuichi Nishinakamura. The murine homolog of SALL4, a causative gene in Okihiro syndrome, is essential for embryonic stem cell proliferation, and cooperates with Sall1 in anorectal, heart, brain and kidney development. Development, 133(15):3005–3013, 2006. ISSN 0950-1991, 1477-9129. doi: 10.1242/dev.02457.

Maria Tahara, Hiroko Kawakami, Katherine Q. Chen, Aaron Anderson, Malina Yamashita Peterson, Wuming Gong, Pruthvi Shah, Shinichi Hayashi, Ryuichi Nishinakamura, Yasushi Nakagawa, Daniel J. Garry, and Yasuhiko Kawakami. Sall4 regulates neuromesodermal progenitors and their descendants during body elongation in mouse embryos. Development, 146(14), 2019. ISSN 0950-1991, 1477-9129. doi: 10.1242/dev.177659.

Maria Akiyama, Hiroko Kawakami, Julia Wong, Isao Oishi, Ryuichi Nishinakamura, and Yasuhiko Kawakami. Sall4-Gli3 system in early limb progenitors is essential for the development of limb skeletal elements. Proceedings of the National Academy of Sciences, 112(16):5075–5080, 2015. ISSN 0027-8424, 1091-6490. doi: 10.1073/pnas.1421949112.

Kazuko Koshiba-Takeuchi, Jun K. Takeuchi, Eric P. Arruda, Irfan S. Kathiriya, Rong Mo, Chi-chung Hui, Deepak Srivastava, and Benoit G. Bruneau. Cooperative and antagonistic interactions between Sall4 and Tbx5 pattern the mouse limb and heart. Nature Genetics, 38(2):175, 2006. ISSN 1546-1718. doi: 10.1038/ng1707.

Ai-Leen Chan, Hue M. La, Julien M. D. Legrand, Juho-Antti Mäkelä, Michael Eichenlaub, Mia De Seram, Mirana Ramialison, and Robin M. Hobbs. Germline Stem Cell Activity Is Sustained by SALL4-Dependent Silencing of Distinct Tumor Suppressor Genes. Stem Cell Reports, 9(3):956–971, 2017. ISSN 2213-6711. doi: 10.1016/j.stemcr.2017.08.001.

Robin M. Hobbs, Sharmila Fagoonee, Antonella Papa, Kaitlyn Webster, Fiorella Altruda, Ryuichi Nishinakamura, Li Chai, and Pier Paolo Pandolfi. Functional Antagonism between Sall4 and Plzf Defines Germline Progenitors. Cell Stem Cell, 10(3):284–298, 2012. ISSN 1934-5909. doi: 10.1016/j.stem.2012.02.004.

Maria Xu, Xia Chen, Hui Yang, Yiwen Xu, Yuanlin He, Chenfei Wang, Hua Huang, Baodong Liu, Wenqiang Liu, Jingyi Li, Xiaochen Kou, Yanhong Zhao, Kun Zhao, Linfeng Zhang, Zhenzhen Hou, Hong Wang, Hailin Wang, Jing Li, Hengyu Fan, Fengchao Wang, Yawei Gao, Yong Zhang, Jiayu Chen, and Shaorong Gao. Maternal Sall4 Is Indispensable for Epigenetic Maturation of Mouse Oocytes. Journal of Biological Chemistry, 292(5):1798–1807, 2017. ISSN 0021-9258, 1083-351X. doi: 10.1074/jbc.M116.767061.

Yasuka L. Yamaguchi, Satomi S. Tanaka, Maho Kumagai, Yuka Fujimoto, Takeshi Terabayashi, Yasuhisa Matsui, and Ryuichi Nishinakamura. Sall4 Is Essential for Mouse Primordial Germ Cell Specification by Suppressing Somatic Cell Program Genes. STEM CELLS, 33(1):289–300, 2015. ISSN 1549-4918 doi: 10.1002/stem.1853.

Maria Elling, Christian Klasen, Tobias Eisenberger, Katrin Anlag, and Mathias Treier. Murine inner cell mass-derived lineages depend on Sall4 function. Proceedings of the National Academy of Sciences, 103(44):16319–16324, 2006. ISSN 0027-8424, 1091-6490. doi: 10.1073/pnas.0607884103.

Maria Warren, Wei Wang, Sarah Spiden, Dongrong Chen-Murchie, David Tannahill, Karen P. Steel, and Allan Bradley. A Sall4 mutant mouse model useful for studying the role of Sall4 in early embryonic development and organogenesis. genesis, 45(1): 51–58, 2007. ISSN 1526-968X. doi: 10.1002/dvg.20264.

Raidah Al-Baradie, Koki Yamada, Cynthia St. Hilaire, Wai-Man Chan, Caroline Andrews, Nathalie McIntosh, Motoi Nakano, E. Jean Martonyi, William R. Raymond, Sada Okumura, Michael M. Okihiro, and Elizabeth C. Engle. Duane Radial Ray Syndrome (Okihiro Syndrome) Maps to 20q13 and Results from Mutations in SALL4, a New Member of the SAL Family. The American Journal of Human Genetics, 71(5):1195–1199, 2002. ISSN 0002-9297. doi: 10.1086/343821.

Jürgen Kohlhase, Marielle Heinrich, Lucia Schubert, Manuela Liebers, Andreas Kispert, Franco Laccone, Peter Turnpenny, Robin M. Winter, and William Reardon. Okihiro syndrome is caused by SALL4 mutations. Human Molecular Genetics, 11(23):2979–2987, 2002. ISSN 0964-6906. doi: 10.1093/hmg/11.23.2979.

Katherine A Donovan, Jian An, Radosław P Nowak, Jingting C Yuan, Emma C Fink, Bethany C Berry, Benjamin L Ebert, and Eric S Fischer. Thalidomide promotes degradation of SALL4, a transcription factor implicated in Duane Radial Ray syndrome. eLife, 7: e38430, 2018. ISSN 2050-084X. doi: 10.7554/eLife.38430.

Mary E. Matyskiela, Suzana Couto, Xinde Zheng, Gang Lu, Julia Hui, Katie Stamp, Clifton Drew, Yan Ren, Maria Wang, Aaron Carpenter, Chung-Wein Lee, Thomas Clayton, Wei Fang, Chin-Chun Lu, Mariko Riley, Polat Abdubek, Kate Blease, James Hartke, Gondi Kumar, Rupert Vessey, Mark Rolfe, Lawrence G. Hamann, and Philip P. Chamberlain. SALL4 mediates teratogenicity as a thalidomide-dependent cereblon substrate. Nature Chemical Biology, 14(10):981–987, 2018. ISSN 1552-4469. doi: 10.1038/s41589-018-0129-x.

Shannon M. Lauberth and Michael Rauchman. A Conserved 12-Amino Acid Motif in Sall1 Recruits the Nucleosome Remodeling and Deacetylase Corepressor Complex. Journal of Biological Chemistry, 281(33):23922–23931, 2006. ISSN 0021-9258, 1083-351X. doi: 10.1074/jbc.M513461200.

Maria Lu, Hawon Jeong, Nikki Kong, Youyang Yang, John Carroll, Hongbo R. Luo, Leslie E. Silberstein, YupoMa, and Li Chai. Stem Cell Factor SALL4 Represses the Transcriptions of PTEN and SALL1 through an Epigenetic Repressor Complex. PLOS ONE, 4 (5):e5577, 2009. ISSN 1932-6203. doi: 10.1371/journal.pone.0005577.

Maria Xiong, Zhuqiang Zhang, Jiayu Chen, Hua Huang, Yali Xu, Xiaojun Ding, Yong Zheng, Ryuichi Nishinakamura, Guo-Liang Xu, Hailin Wang, She Chen, Shaorong Gao, and Bing Zhu. Cooperative Action between SALL4A and TET Proteins in Stepwise Oxidation of 5-Methylcytosine. Molecular Cell, 64(5):913–925, 2016. ISSN 10972765. doi: 10.1016/j.molcel.2016.10.013.

Maria Tanimura, Motoki Saito, Miki Ebisuya, Eisuke Nishida, and Fuyuki Ishikawa. Stemness-related Factor Sall4 Interacts with Transcription Factors Oct-3/4 and Sox2 and Occupies Oct-Sox Elements in Mouse Embryonic Stem Cells. Journal of Biological Chemistry, 288(7):5027–5038, 2013. ISSN 0021-9258, 1083-351X. doi: 10.1074/jbc.M112.411173.

Cornelia G. Spruijt, H. Irem Baymaz, and Michiel Vermeulen. Identifying Specific Protein–DNA Interactions Using SILAC-Based Quantitative Proteomics. In Minou Bina, editor, Gene Regulation: Methods and Protocols, Methods in Molecular Biology, pages 137–157. Humana Press, Totowa, NJ, 2013a. ISBN 978-1-62703-284-1.

Cornelia G. Spruijt, Felix Gnerlich, Arne H. Smits, Toni Pfaffeneder, Pascal W.T.C. Jansen, Christina Bauer, Martin Münzel, Mirko Wagner, Markus Müller, Fariha Khan, H. Christian Eberl, Anneloes Mensinga, Arie B. Brinkman, Konstantin Lephikov, Udo Müller, Jörn Walter, Rolf Boelens, Hugo van Ingen, Heinrich Leonhardt, Thomas Carell, and Michiel Vermeulen. Dynamic Readers for 5-(Hydroxy)Methylcytosine and Its Oxidized Derivatives. Cell, 152(5):1146–1159, 2013b. ISSN 00928674. doi: 10.1016/j.cell.2013.02.004.

L. Aravind and David Landsman. AT-hook motifs identified in a wide variety of DNA-binding proteins. Nucleic Acids Research, 26 (19):4413–4421, 1998. ISSN 0305-1048. doi: 10.1093/nar/26.19.4413.

Maria Filarsky, Karina Zillner, Ingrid Araya, Ana Villar-Garea, Rainer Merkl, Gernot Längst, and Attila Németh. The extended AT-hook is a novel RNA binding motif. RNA Biology, 12(8):864–876, 2015. ISSN 1547-6286. doi: 10.1080/15476286.2015.1060394.

Maria Patsialou, Rosalia Wilsker, and Elizabeth Moran. DNA-binding properties of ARID family proteins. Nucleic Acids Research, 33(1):66–80, 2005. ISSN 0305-1048. doi: 10.1093/nar/gki145.

Dylan Sweetman and Andrea Münsterberg. The vertebrate spalt genes in development and disease. Developmental Biology, 293(2): 285–293, 2006. ISSN 0012-1606. doi: 10.1016/j.ydbio.2006.02.009.

Jeffrey K. Tong, Christian A. Hassig, Gavin R. Schnitzler, Robert E. Kingston, and Stuart L. Schreiber. Chromatin deacetylation by an ATP-dependent nucleosome remodelling complex. Nature, 395(6705):917, 1998. ISSN 1476-4687. doi: 10.1038/27699.

Paul A. Wade, Peter L. Jones, Danielle Vermaak, and Alan P. Wolffe. A multiple subunit Mi-2 histone deacetylase from Xenopus laevis cofractionates with an associated Snf2 superfamily ATPase. Current Biology, 8(14):843–848, 1998. ISSN 0960-9822. doi: 10.1016/S0960-9822(98)70328-8.

Maria Xue, Jiemin Wong, G. Tony Moreno, Mary K. Young, Jacques Côté, and Weidong Wang. NURD, a Novel Complex with Both ATP-Dependent Chromatin-Remodeling and Histone Deacetylase Activities. Molecular Cell, 2(6):851–861, 1998. ISSN 1097-2765. doi: 10.1016/S1097-2765(00)80299-3.

Maria Zhang, Gary LeRoy, Hans-Peter Seelig, William S Lane, and Danny Reinberg. The Dermatomyositis-Specific Autoantigen Mi2 Is a Component of a Complex Containing Histone Deacetylase and Nucleosome Remodeling Activities. Cell, 95(2):279–289, 1998. ISSN 0092-8674. doi: 10.1016/S0092-8674(00)81758-4.

Maria Yamashita, Akira Sato, Makoto Asashima, Pi-Chao Wang, and Ryuichi Nishinakamura. Mouse homolog of SALL1, a causative gene for Townes–Brocks syndrome, binds to A/T-rich sequences in pericentric heterochromatin via its C-terminal zinc finger domains. Genes to Cells, 12(2):171–182, 2007. ISSN 1365-2443. doi: 10.1111/j.1365-2443.2007.01042.x.

Yoichi Matsuda and Verne M. Chapman. In situ analysis of centromeric satellite DNA segregating inMus species crosses. Mammalian Genome, 1(2):71, 1991. ISSN 1432-1777. doi: 10.1007/BF02443781.

María Cristina Cerda, Soledad Berríos, Raúl Fernández-Donoso, Silvia Garagna, and Carlo Redi. Organisation of complex nuclear domains in somatic mouse cells. Biology of the Cell, 91(1):55–65, 1999. ISSN 1768-322X. doi: 10.1111/j.1768-322X.1999.tb01084.x.

Maria Guenatri, Delphine Bailly, Christèle Maison, and Geneviève Almouzni. Mouse centric and pericentric satellite repeats form distinct functional heterochromatin. Journal of Cell Biology, 166(4):493–505, 2004. ISSN 0021-9525. doi: 10.1083/jcb.200403109.

Maria Jolma, Teemu Kivioja, Jarkko Toivonen, Lu Cheng, Gonghong Wei, Martin Enge, Mikko Taipale, Juan M. Vaquerizas, Jian Yan, Mikko J. Sillanpää, Martin Bonke, Kimmo Palin, Shaheynoor Talukder, Timothy R. Hughes, Nicholas M. Luscombe, Esko Ukkonen, and Jussi Taipale. Multiplexed massively parallel SELEX for characterization of human transcription factor binding specificities. Genome Research, 20(6):861–873, 2010. ISSN 1088-9051, 1549-5469. doi: 10.1101/gr.100552.109.

Kazuhiro R Nitta, Arttu Jolma, Yimeng Yin, Ekaterina Morgunova, Teemu Kivioja, Junaid Akhtar, Korneel Hens, Jarkko Toivonen, Bart Deplancke, Eileen E M Furlong, and Jussi Taipale. Conservation of transcription factor binding specificities across 600 million years of bilateria evolution. eLife, 4:e04837, 2015. ISSN 2050-084X. doi: 10.7554/eLife.04837.

Benjamin L. Kidder, Gangqing Hu, and Keji Zhao. ChIP-Seq: technical considerations for obtaining high-quality data. Nature Immunology, 12(10):918–922, 2011. ISSN 1529-2916. doi: 10.1038/ni.2117.

Stephen G. Landt, Georgi K. Marinov, Anshul Kundaje, Pouya Kheradpour, Florencia Pauli, Serafim Batzoglou, Bradley E. Bernstein, Peter Bickel, James B. Brown, Philip Cayting, Yiwen Chen, Gilberto DeSalvo, Charles Epstein, Katherine I. Fisher-Aylor, Ghia Euskirchen, Mark Gerstein, Jason Gertz, Alexander J. Hartemink, Michael M. Hoffman, Vishwanath R. Iyer, Youngsook L. Jung, Subhradip Karmakar, Manolis Kellis, Peter V. Kharchenko, Qunhua Li, Tao Liu, X. Shirley Liu, Lijia Ma, Aleksandar Milosavljevic, Richard M. Myers, Peter J. Park, Michael J. Pazin, Marc D. Perry, Debasish Raha, Timothy E. Reddy, Joel Rozowsky, Noam Shoresh, Arend Sidow, Matthew Slattery, John A. Stamatoyannopoulos, Michael Y. Tolstorukov, Kevin P. White, Simon Xi, Peggy J. Farnham, Jason D. Lieb, Barbara J. Wold, and Michael Snyder. ChIP-seq guidelines and practices of the ENCODE and modENCODE consortia. Genome Research, 22(9):1813–1831, 2012. ISSN 1088-9051, 1549-5469. doi: 10.1101/gr.136184.111.

Maria Uhlen, Anita Bandrowski, Steven Carr, Aled Edwards, Jan Ellenberg, Emma Lundberg, David L. Rimm, Henry Rodriguez, Tara Hiltke, Michael Snyder, and Tadashi Yamamoto. A proposal for validation of antibodies. Nature Methods, 13(10):823–827, 2016. ISSN 1548-7105. doi: 10.1038/nmeth.3995.

Maria Chronis, Petko Fiziev, Bernadett Papp, Stefan Butz, Giancarlo Bonora, Shan Sabri, Jason Ernst, and Kathrin Plath. Cooperative Binding of Transcription Factors Orchestrates Reprogramming. Cell, 168(3):442–459.e20, January 2017. ISSN 0092-8674. doi: 10.1016/j.cell.2016.12.016.

Maria Rao, Shao Zhen, Sergei Roumiantsev, Lindsay T. McDonald, Guo-Cheng Yuan, and Stuart H. Orkin. Differential Roles of Sall4 Isoforms in Embryonic Stem Cell Pluripotency. Molecular and Cellular Biology, 30(22):5364–5380, 2010. ISSN 0270-7306, 1098-5549. doi: 10.1128/MCB.00419-10.

Jerôme Aubert, Hannah Dunstan, Ian Chambers, and Austin Smith. Functional gene screening in embryonic stem cells implicates Wnt antagonism in neural differentiation. Nature Biotechnology, 20(12):1240–1245, 2002. ISSN 1087-0156. doi: 10.1038/nbt763.

Maria Tsubooka, Tomoko Ichisaka, Keisuke Okita, Kazutoshi Takahashi, Masato Nakagawa, and Shinya Yamanaka. Roles of Sall4 in the generation of pluripotent stem cells from blastocysts and fibroblasts. Genes to Cells, 14(6):683–694, 2009. ISSN 1365-2443. doi: 10.1111/j.1365-2443.2009.01301.x.

Maria Terhal, Bernd Rösler, and Jürgen Kohlhase. A family with features overlapping Okihiro syndrome, hemifacial microsomia and isolated Duane anomaly caused by a novel SALL4 mutation. American Journal of Medical Genetics Part A, 140A(3):222–226, 2006. ISSN 1552-4833. doi: 10.1002/ajmg.a.31060.

Maria Miertus, Wiktor Borozdin, Vladimir Frecer, Giorgio Tonini, Sara Bertok, Antonio Amoroso, Stanislav Miertus, and Jürgen Kohlhase. A SALL4 zinc finger missense mutation predicted to result in increased DNA binding affinity is associated with cranial midline defects and mild features of Okihiro syndrome. Human Genetics, 119(1-2):154–161, 2006. ISSN 0340-6717, 1432-1203. doi: 10.1007/s00439-005-0124-7.

Maria Zhang, Arne H. Smits, Gabrielle B. A. van Tilburg, Pascal W. T. C. Jansen, Matthew M. Makowski, Huib Ovaa, and Michiel Vermeulen. An Interaction Landscape of Ubiquitin Signaling. Molecular Cell, 65(5):941–955.e8, 2017. ISSN 1097-2765. doi: 10.1016/j.molcel.2017.01.004.

Susan McLeskey Kiefer, Kevin K. Ohlemiller, Jing Yang, Bradley W. McDill, Jürgen Kohlhase, and Michael Rauchman. Expression of a truncated Sall1 transcriptional repressor is responsible for Townes–Brocks syndrome birth defects. Human Molecular Genetics, 12(17):2221–2227, 2003. ISSN 0964-6906. doi: 10.1093/hmg/ddg233.

Maria Sweetman, Terry Smith, Elizabeth R. Farrell, Andrew Chantry, and Andrea Münsterberg. The Conserved Glutamine-rich Region of Chick Csal1 and Csal3 Mediates Protein Interactions with Other Spalt Family Members IMPLICATIONS FOR TOWNES-BROCKS SYNDROME. Journal of Biological Chemistry, 278(8):6560–6566, 2003. ISSN 0021-9258, 1083-351X. doi: 10.1074/jbc.M209066200.

Maria Hooper, Kate Hardy, Alan Handyside, Susan Hunter, and Marilyn Monk. HPRT-deficient (Lesch–Nyhan) mouse embryos derived from germline colonization by cultured cells. Nature, 326(6110):292–295, 1987. ISSN 0028-0836. doi: 10.1038/326292a0.

Qi-Long Ying, Jason Wray, Jennifer Nichols, Laura Batlle-Morera, Bradley Doble, James Woodgett, Philip Cohen, and Austin Smith. The ground state of embryonic stem cell self-renewal. Nature, 453(7194):519–523, 2008. ISSN 0028-0836. doi: 10.1038/nature06968.

F. Ann Ran, Patrick D. Hsu, Jason Wright, Vineeta Agarwala, David A. Scott, and Feng Zhang. Genome engineering using the CRISPR-Cas9 system. Nature Protocols, 8(11):2281–2308, 2013. ISSN 1754-2189. doi: 10.1038/nprot.2013.143.

Maria Yusa, Liqin Zhou, Meng Amy Li, Allan Bradley, and Nancy L. Craig. A hyperactive piggyBac transposase for mammalian applications. Proceedings of the National Academy of Sciences, 108(4):1531–1536, 2011. ISSN 0027-8424, 1091-6490. doi: 10.1073/pnas.1008322108.

Maria Rappsilber, Rosalia Ishihama, and Matthias Mann. Stop and Go Extraction Tips for Matrix-Assisted Laser Desorption/Ionization, Nanoelectrospray, and LC/MS Sample Pretreatment in Proteomics. Analytical Chemistry, 75(3):663–670, 2003. ISSN 0003-2700. doi: 10.1021/ac026117i.

Jürgen Cox and Matthias Mann. MaxQuant enables high peptide identification rates, individualized p.p.b.-range mass accuracies and proteome-wide protein quantification. Nature Biotechnology, 26(12):1367–1372, 2008. ISSN 1546-1696. doi: 10.1038/nbt.1511.

Maria Tyanova, Tikira Temu, Pavel Sinitcyn, Arthur Carlson, Marco Y. Hein, Tamar Geiger, Matthias Mann, and Jürgen Cox. The Perseus computational platform for comprehensive analysis of (prote)omics data. Nature Methods, 13(9):731–740, 2016. ISSN 1548-7105. doi: 10.1038/nmeth.3901.

Julie K. Stock, Sara Giadrossi, Miguel Casanova, Emily Brookes, Miguel Vidal, Haruhiko Koseki, Neil Brockdorff, Amanda G. Fisher, and Ana Pombo. Ring1-mediated ubiquitination of H2A restrains poised RNA polymerase II at bivalent genes in mouse ES cells. Nature Cell Biology, 9(12):1428–1435, 2007. ISSN 1476-4679. doi: 10.1038/ncb1663.

Justyna Cholewa-Waclaw, Ruth Shah, Shaun Webb, Kashyap Chhatbar, Bernard Ramsahoye, Oliver Pusch, Miao Yu, Philip Greulich, Bartlomiej Waclaw, and Adrian P. Bird. Quantitative modelling predicts the impact of DNA methylation on RNA polymerase II traffic. Proceedings of the National Academy of Sciences, 116(30):14995–15000, 2019. ISSN 0027-8424, 1091-6490. doi: 10.1073/pnas.1903549116.

Kushal K. Dey, Dongyue Xie, and Matthew Stephens. A new sequence logo plot to highlight enrichment and depletion. BMC Bioinformatics, 19(1):473, 2018. ISSN 1471-2105. doi: 10.1186/s12859-018-2489-3.

Maria Patro, Stephen M. Mount, and Carl Kingsford. Sailfish enables alignment-free isoform quantification from RNA-seq reads using lightweight algorithms. Nature Biotechnology, 32(5):462–464, 2014. ISSN 1546-1696. doi: 10.1038/nbt.2862.

Michael I. Love, Wolfgang Huber, and Simon Anders. Moderated estimation of fold change and dispersion for RNA-seq data with DESeq2. Genome Biology, 15(12):550, 2014. ISSN 1474-760X. doi: 10.1186/s13059-014-0550-8.

Aaron R. Quinlan and Ira M. Hall. BEDTools: a flexible suite of utilities for comparing genomic features. Bioinformatics, 26(6): 841–842, 2010. ISSN 1367-4803. doi: 10.1093/bioinformatics/btq033.

Fidel Ramírez, Friederike Dündar, Sarah Diehl, Björn A. Grüning, and Thomas Manke. deepTools: a flexible platform for exploring deep-sequencing data. Nucleic Acids Research, 42(W1):W187–W191, 2014. ISSN 0305-1048. doi: 10.1093/nar/gku365.

Maria Yu, Li-Gen Wang, Yanyan Han, and Qing-Yu He. clusterProfiler: an R Package for Comparing Biological Themes Among Gene Clusters. OMICS: A Journal of Integrative Biology, 16(5):284–287, 2012. doi: 10.1089/omi.2011.0118.

Anthony M. Bolger, Marc Lohse, and Bjoern Usadel. Trimmomatic: a flexible trimmer for Illumina sequence data. Bioinformatics, 30(15):2114–2120, 2014. ISSN 1367-4803. doi: 10.1093/bioinformatics/btu170.

Heng Li. Aligning sequence reads, clone sequences and assembly contigs with BWA-MEM. arXiv:1303.3997 [q-bio], 2013.

Yuval Benjamini and Terence P. Speed. Summarizing and correcting the GC content bias in high-throughput sequencing. Nucleic Acids Research, 40(10):e72–e72, 2012. ISSN 0305-1048. doi: 10.1093/nar/gks001.

Maria Zhang, Tao Liu, Clifford A. Meyer, Jérôme Eeckhoute, David S. Johnson, Bradley E. Bernstein, Chad Nusbaum, Richard M. Myers, Myles Brown, Wei Li, and X. Shirley Liu. Model-based Analysis of ChIP-Seq (MACS). Genome Biology, 9:R137, 2008. ISSN 1474-760X. doi: 10.1186/gb-2008-9-9-r137.

Philip Machanick and Timothy L. Bailey. MEME-ChIP: motif analysis of large DNA datasets. Bioinformatics, 27(12):1696–1697, 2011. ISSN 1367-4803. doi: 10.1093/bioinformatics/btr189.

