## Supplementary Figures and Bioinformatics analyses for "SALL4 controls cell fate in response to DNA base composition"

#### Contents

**Figure S1** 2

**Figure S2** 3

**Figure S3** 4

**Figure S4** 6

**Figure S5** 6

**Figure S6** 7

**Figure S7** 8

**Bioinformatics Analyses** 9

[*]These authors contributed equally
[†]The Wellcome Centre for Cell Biology, University of Edinburgh, Michael Swann Building, Max Born Crescent, The King's Buildings, Edinburgh, EH9 3BF, UK
[‡]Informatics Forum, School of Informatics, University of Edinburgh, 10 Crichton Street, Edinburgh, EH8 9AB, UK
[§]Department of Molecular Biology, Faculty of Science, Radboud Institute for Molecular Life Sciences, Oncode Institute, Radboud University Nijmegen, Nijmegen, The Netherlands
[||]Corresponding author

#### Figure S1

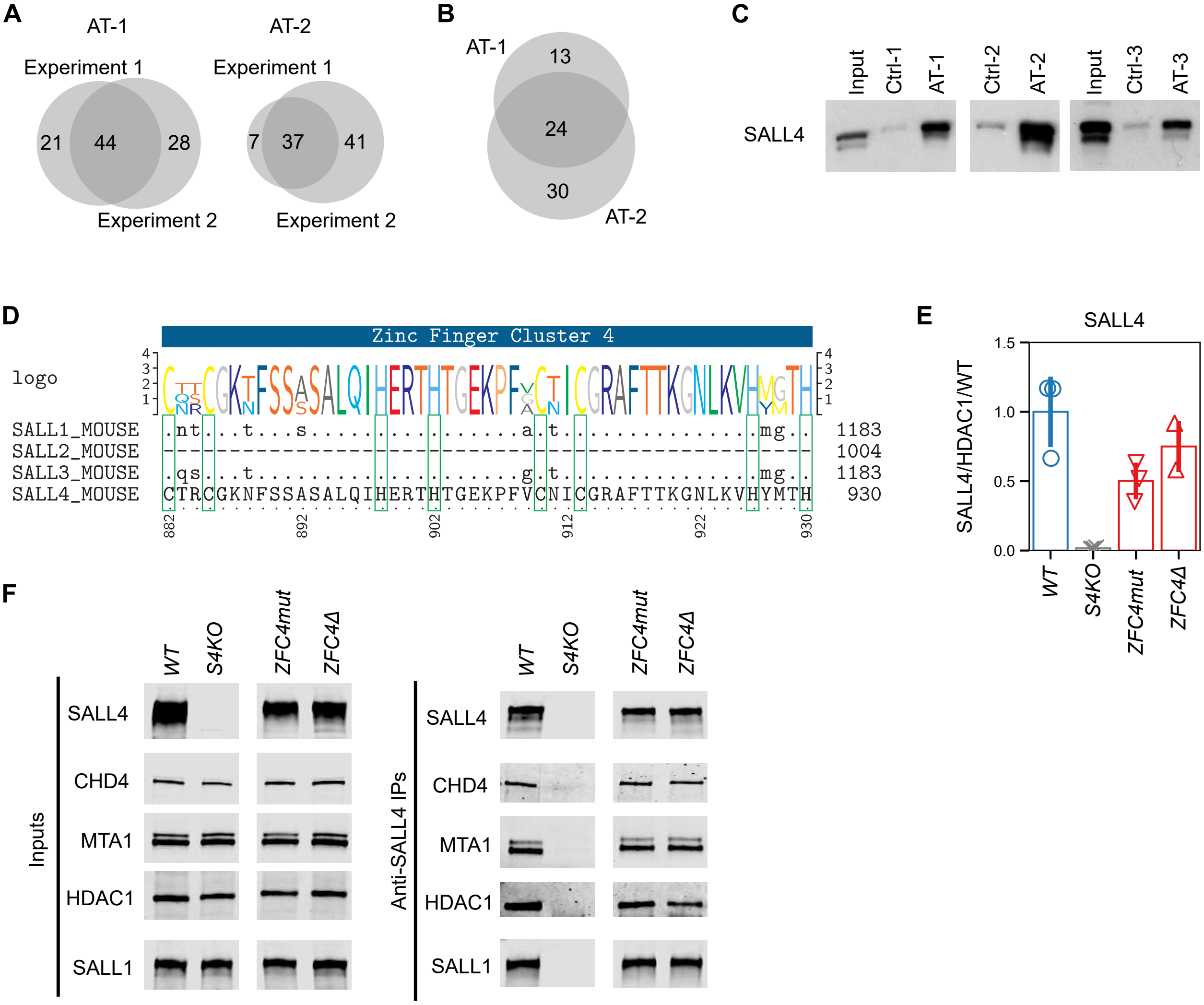

**Supplemental Figure 1 (related to Figure 1)**

**A, B.** Venn diagrams showing the overlap between proteins identified by DNA pulldown-mass spectrometry in independent replicate experiments (A), or using unrelated AT-rich DNA probes (B). **C.** DNA pulldown with AT-rich (AT-1, AT-2, AT-3) or control (Ctrl-1, Ctrl-2, Ctrl-3) probes followed by Western blot analysis for SALL4 using *WT* ESC protein extracts. **D.** Protein alignment and consensus sequence of C2H2 zinc-finger cluster 4 (ZFC4) in the mouse SALL protein family. ZFC4 is absent in SALL2. **E.** Western blot quantification of SALL4 expression levels in *S4KO* and *ZFC4mut*/Δ ESCs, normalised to HDAC1 expression and relative to *WT* ESC levels. Data points indicate independent replicate experiments and error bars standard deviation. **F.** SALL4 co-immunoprecipitation with SALL1 and NuRD components in *WT*, *S4KO* (negative control) and *ZFC4mut*/Δ ESCs. For both inputs and anti-SALL4 IPs, all four lanes are part of the same Western blot membrane and images were processed in an identical manner.

#### Figure S2

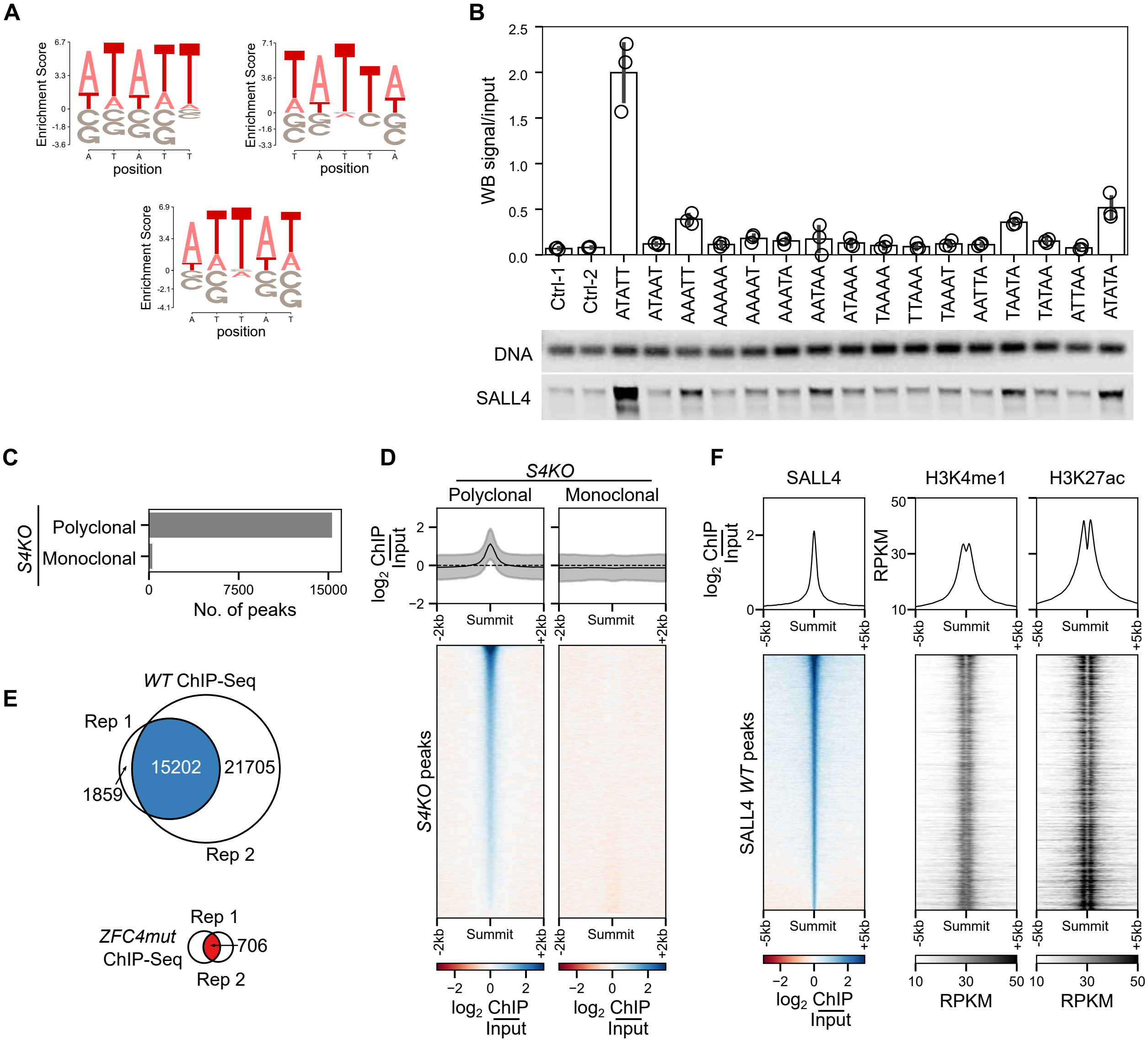

**Supplemental Figure 2 (related to Figure 2)**
**A.** Motif logos generated from the three most enriched k-mers (n=5) after 6 cycles of HT-SELEX with SALL4 ZFC4. **B.** DNA pulldown with AT-rich probes containing all possible combinations of AT 5 mers or control probes with disrupted AT-runs (Ctrl-) followed by Western blot analysis for SALL4. Amounts of DNA probes were assessed by agarose gel analysis and SALL4 enrichment was normalised to input. Data points indicate independent replicate experiments and error bars standard deviation. **C.** Detection of non-specific SALL4 ChIP-seq peaks in *Sall4* knockout ESCs (negative control) using either a monoclonal or a polyclonal anti-SALL4 antibody. **D.** Profile plot and heatmap showing SALL4 ChIP-seq signal in *Sall4* knockout ESCs at non-specific sites (see panel B) using either a monoclonal or a polyclonal anti-SALL4 antibody. **E.** Venn diagrams showing the overlap of SALL4 ChIP-seq peaks between independent replicate experiments using an anti-SALL4 monoclonal antibody in *WT* (blue) and *ZFC4mut* (red) ESC lines. **F.** Profile plots and heatmaps showing SALL4, H3K4me1 and H3K27ac ChIP-seq signal at SALL4 *WT* ChIP-seq peaks in *WT* ESCs.

### Figure S3

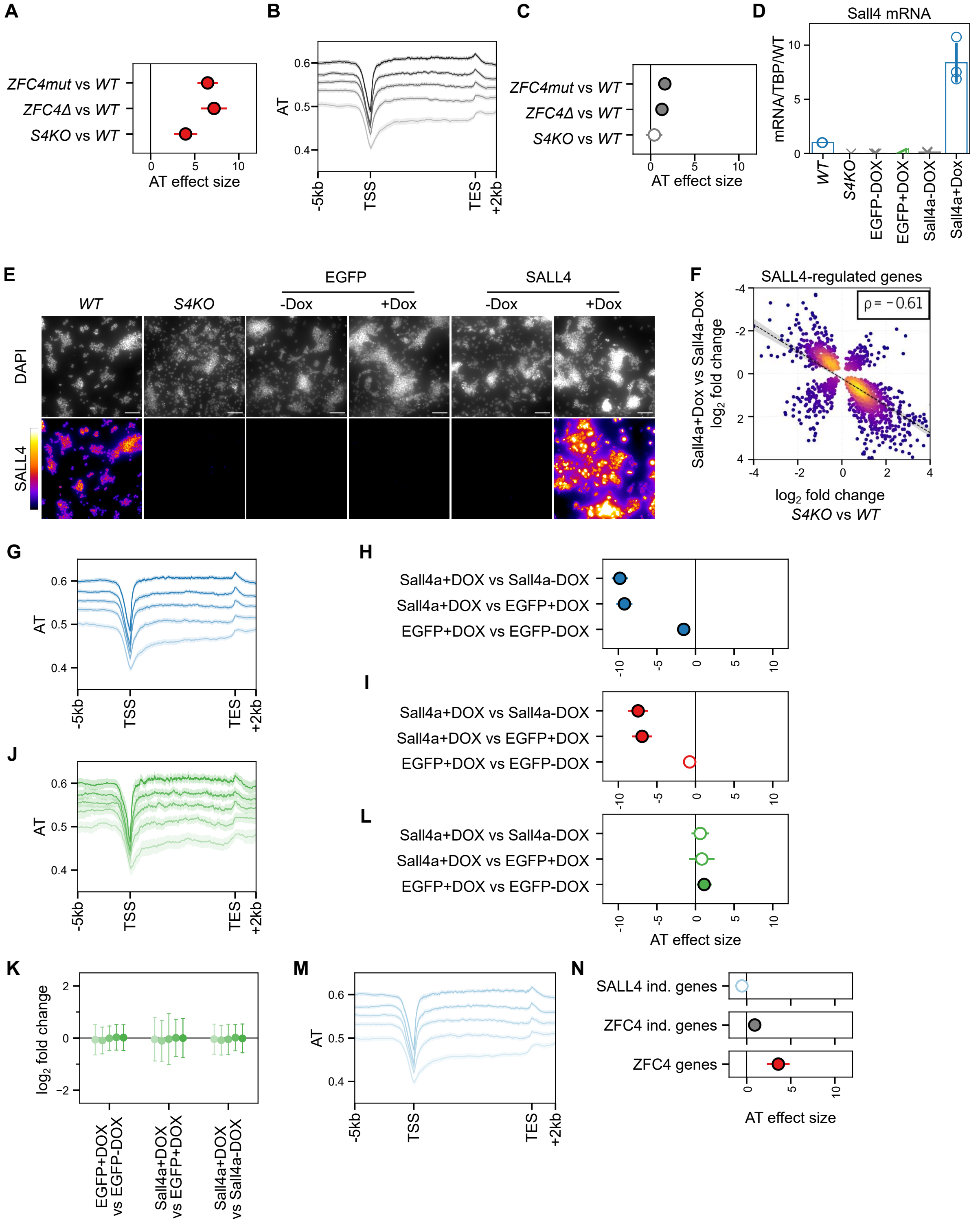

**Supplemental Figure 3 (related to Figure 3)**

*(continued)*

**Supplemental Figure 3 (related to Figure 3):** ***(continued)***

**A.** Statistical analysis of AT-dependent gene expression changes (coefficient estimates with 99% confidence intervals) observed with ZFC4-regulated genes (see Figure 3A). Significance is attributed by the F-test. Empty circles represent non-significant model fits (>0.01 FDR) and filled circles represent a significant fit to the model. **B.** Profile plot showing the density of A/T nucleotides around the transcription unit of ZFC4-independent genes (see Figure 3A) divided into five equal categories according to AT-content. **C.** Statistical analysis of AT-dependent gene expression changes observed with ZFC4-independent genes, as described in panel A. **D.** RT-qPCR analysis following 48h doxycycline induction in the indicated ESC lines (see Figure 3E), or in *WT* and *S4KO* control ESCs. Sall4 mRNA expression was normalised to TBP and expressed relative to *WT*. Data points indicate independent replicate experiments and error bars standard deviation. **E.** SALL4 immunofluorescence following 48h doxycycline induction in the indicated ESC lines (see Figure 3E), or in *WT* and *S4KO* control ESCs. DNA was stained with DAPI. Scale bars: 100µm. **F.** Scatter plot showing the relative expression of genes deregulated both in *S4KO* ESCs and following SALL4 re-expression. **G.** Profile plot showing the density of A/T nucleotides around the transcription unit of Sall4-responsive genes (see Figure 3F) divided into five equal categories according to AT-content. **H, I.** Statistical analysis of AT-dependent gene expression changes observed with Sall4-responsive (H) and ZFC4-regulated (I) genes, as described in panel A. **J.** Profile plot showing the density of A/T nucleotides around the transcription unit of EGFP-responsive genes (see Figure 3F) divided into five equal categories according to AT-content. **K.** Correlation between EGFP-induced gene expression changes and DNA base composition. EGFP-responsive genes were divided into five equal categories depending on their AT-content, and their relative expression levels were analysed in the indicated ESC lines. **L.** Statistical analysis of AT-dependent gene expression changes observed with EGFP-responsive genes, as described in panel A. **M.** Profile plot showing the density of A/T nucleotides around the transcription unit of SALL4-independent genes changing during early ESC differentiation (see Figure 3J) divided into five equal categories according to AT-content. **N.** Statistical analysis of AT-dependent gene expression changes observed with SALL4-independent genes (light blue), SALL4-dependent genes controlled by ZFC4 (red) and SALL4-dependent genes not controlled by ZFC4 (grey) during early differentiation of *WT* cells (day 0 *vs* day 2), as described in panel A.

#### Figure S4

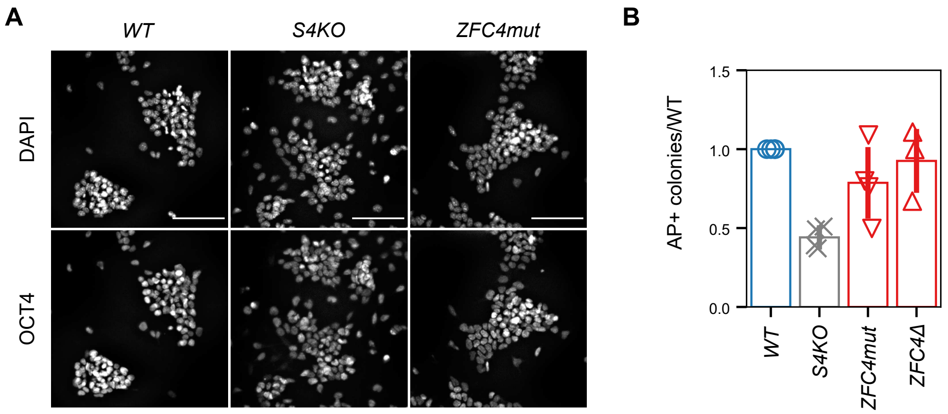

**Supplemental Figure 4 (related to Figure 4)**
**A.** OCT4 immunofluorescence in *WT*, *S4KO* and *ZFC4mut* ESCs. DNA was stained with DAPI. Scale bars: 100μm. **B.** Self-renewal assay in *WT*, *S4KO* and *ZFC4mut*/Δ ESCs. Alkaline phosphatase (AP)-positive colonies were counted and normalised to *WT*. Data points indicate independent replicate experiments and error bars standard deviation.

#### Figure S5

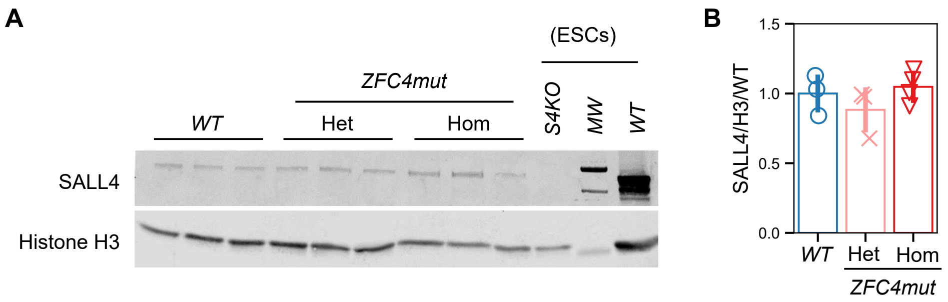

**Supplemental Figure 5 (related to Figure 5)**
**A.** Western blot analysis of SALL4 in *WT*, *ZFC4mut* heterozygote (*Het*) and homozygote (*Hom*) embryos at E10.5. *WT* and *S4KO* ESC protein extracts were used as controls. **B.** Western blot quantification of SALL4 expression levels in *ZFC4mut* embryos (as presented in panel A), normalised to Histone H3 expression and relative to *WT*. Data points indicate independent embryos and error bars standard deviation.

#### Figure S6

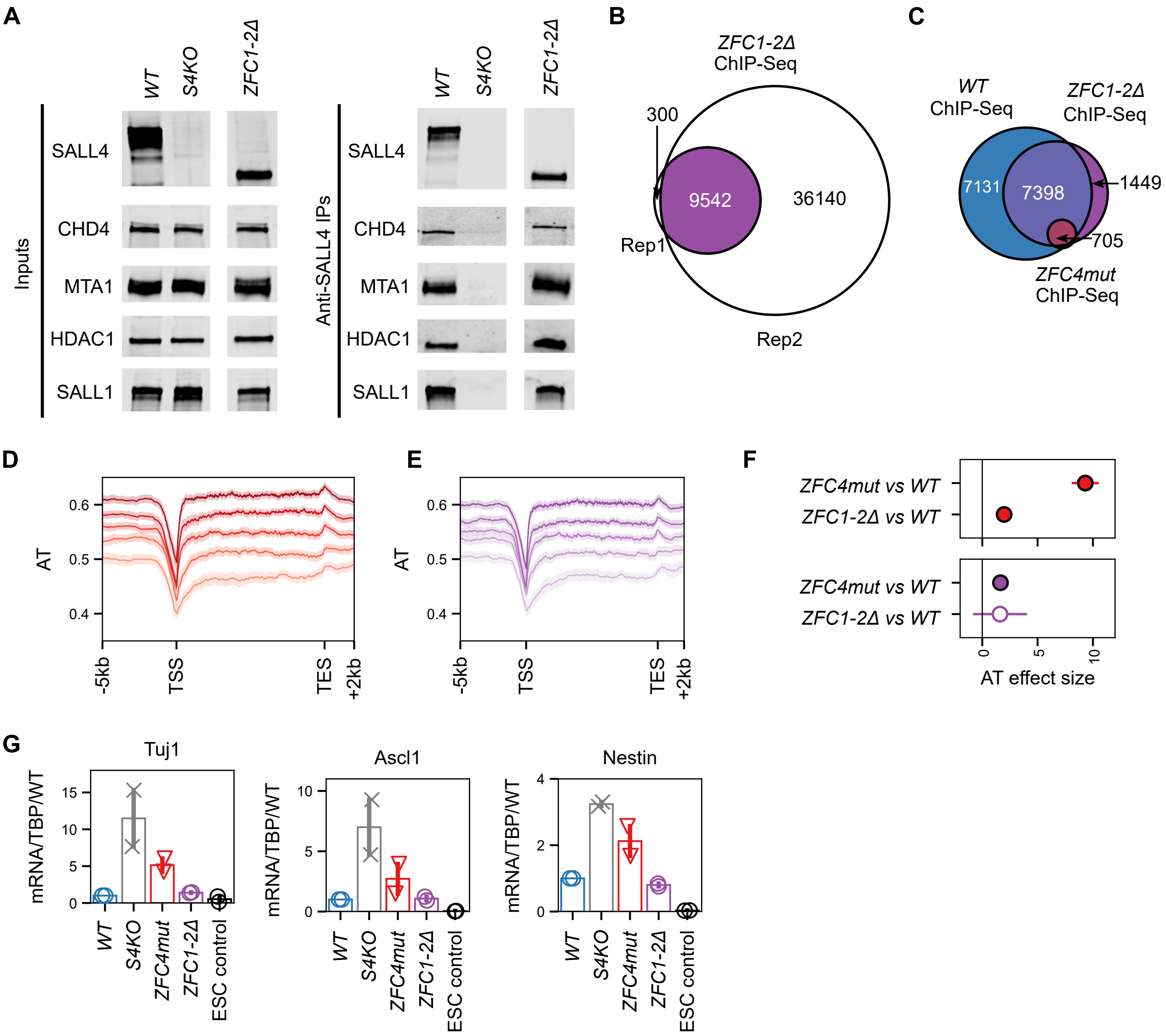

**Supplemental Figure 6 (related to Figure 6)**

**A.** SALL4 co-immunoprecipitation with SALL1 and NuRD components in *WT*, *S4KO* (negative control) and *ZFC1-2*Δ ESCs. For both inputs and anti-SALL4 IPs, all three lanes are part of the same Western blot membrane and images were processed in an identical manner. **B.** Venn diagram showing the overlap of SALL4 ChIP-seq peaks between independent replicate experiments in *ZFC1-2*Δ ESCs. **C.** Venn diagram showing the overlap of SALL4 ChIP-seq peaks between *WT*, *ZFC1-2*Δ and *ZFC4mut* ESCs. **D, E.** Profile plot showing the density of A/T nucleotides around the transcription unit of ZFC4-regulated (D) and ZFC1/2-regulated (E) genes (see Figure 6E) divided into five equal categories according to AT-content. **F.** Statistical analysis of AT-dependent gene expression changes (coefficient estimates with 99% confidence intervals) observed with ZFC4-regulated (red) and ZFC1/2-regulated (purple) genes (see Figure 6E). Significance is attributed by F-test. Empty circles represent non-significant model fits (>0.01 FDR) and filled circles represent significant model fit. **G.** RT-qPCR analysis of the neuronal markers Tuj1, Ascl1 and Nestin in the indicated cell lines following differentiation for 5 days in N2B27 medium. Transcripts levels were normalised to TBP and expressed relative to *WT*. Data points indicate independent replicate experiments and error bars standard deviation.

#### Figure S7

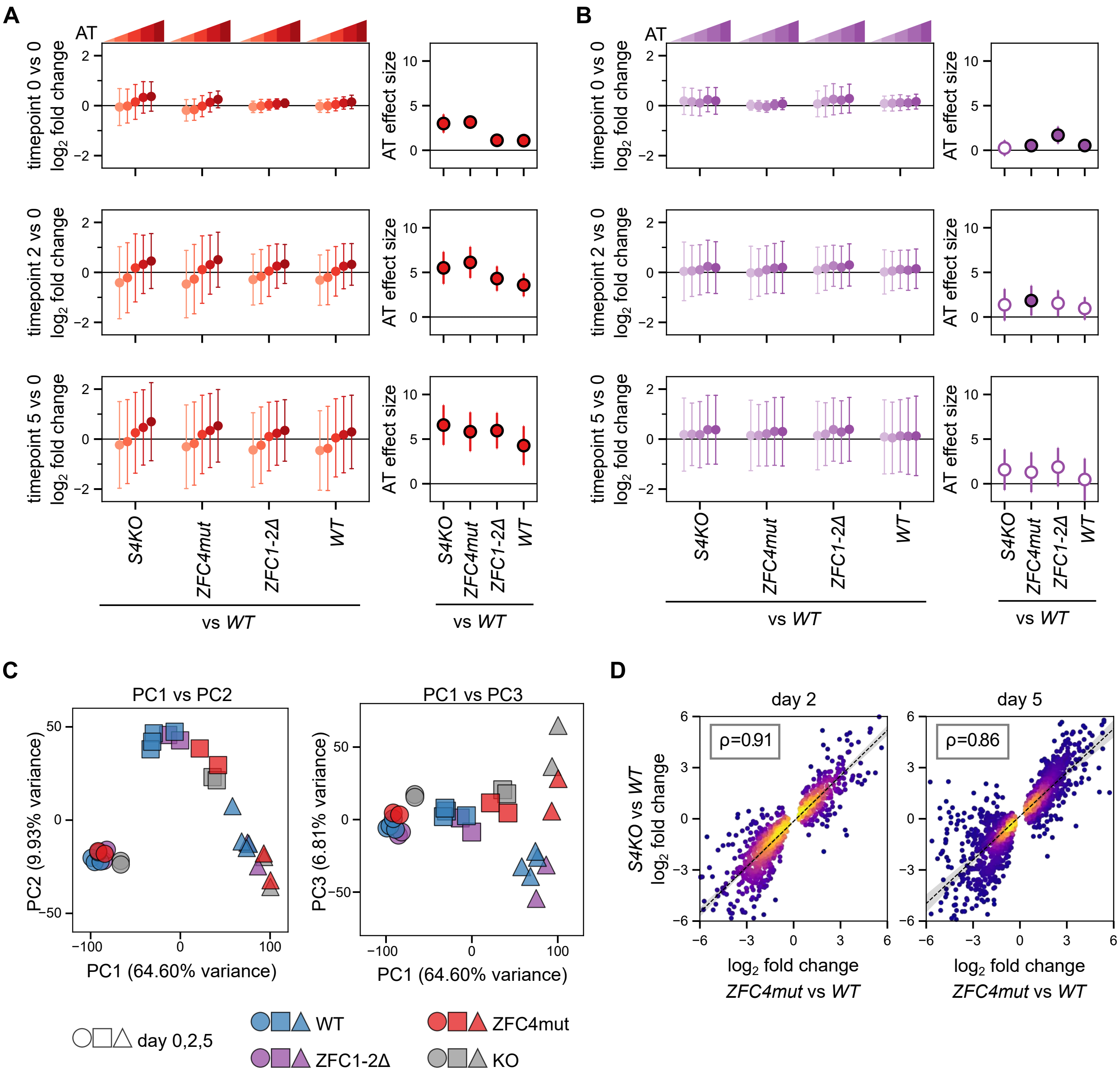

**Supplemental Figure 7 (related to Figure 7)**

**A.** Correlation between gene expression changes and DNA base composition observed with ZFC4-regulated genes at day 0 (top panel), day 2 (middle panel) and day 5 (bottom panel) of differentiation. ZFC4-regulated genes (see Figure 6E) were divided into five equal categories according to their AT-content. Left panel: relative expression levels (log2 fold-change *vs* day 0 in *WT* cells) in *WT* and *Sall4* mutant cells. Right panel: Coefficient estimates (with 99% confidence intervals) describing the AT effect size. **B.** Correlation between gene expression changes and DNA base composition observed with ZFC1/2-regulated genes during differentiation, as described in panel A. **C.** PCA analysis of RNA-seq samples from *WT* and *Sall4* mutant cell lines at day 0, 2 and 5 of differentiation. **D.** Scatter plot showing the relative expression levels of genes deregulated in differentiating *ZFC4mut* cells (see Figure 7B, red bars) correlating with their expression in *S4KO* cells at day 2 and 5 of differentiation.

#### Bioinformatics Analyses

##### Command line arguments for counting k-mers

k-mer abundance was calculated using the following commands

```
jellyfish count -m 5 -C -t 4 -s 100M -o 5.jf <(zcat sample.fq.gz)

jellyfish dump 5.jf > 5_counts.fa
```

Fraction of reads containing k-mers was calculated after executing `calculate_fraction.py` and `calculate_score.py` scripts on counts obtained using the above steps. Scripts are included in supplemental data.

##### Command-line arguments for ChIP-seq anlaysis

```
trimmomatic SE -threads 16 -summary R1.trimmomatic.log R1.fq R1.trimmed.fq
    ↪ ILLUMINACLIP:adapters/TruSeq-SE-combined.fa:2:30:10 LEADING:3 TRAILING:3
    ↪ SLIDINGWINDOW:4:20 MINLEN:36

bwa mem -t 6 -M mm10 R1.trimmed.fq | samtools view -bT mm10.fa -q 1 -F 4 -F 256 >
    ↪ R1.unsorted.bam

samtools sort -o R1.sorted.bam R1.unsorted.bam

samtools index R1.sorted.bam

picard MarkDuplicates I=R1.sorted.bam O=R1.dedup.sorted.bam ASSUME_SORTED=true
    ↪ REMOVE_DUPLICATES=true METRICS_FILE=R1.dedup.metrics VALIDATION_STRINGENCY=LENIENT
    ↪ PROGRAM_RECORD_ID=’null’

samtools index R1.dedup.sorted.bam

computeGCBias -b R1.dedup.sorted.bam --effectiveGenomeSize 2494787188 -g mm10.2bit -bl
    ↪ blacklist.bed -p 20 -l 240 -o R1.dedup.gcbias.freq --biasPlot R1.dedup.gcbias.png

correctGCBias -b R1.dedup.sorted.bam --effectiveGenomeSize 2494787188 -g mm10.2bit -p 20
    ↪ -freq R1.dedup.gcbias.freq -o R1.dedup.sorted.gc.corrected.bam

samtools index R1.dedup.sorted.gc.corrected.bam

macs2 callpeak -t R1.chip.dedup.sorted.gc.corrected.bam -c
    ↪ R1.control.dedup.sorted.gc.corrected.bam -f BAM -g 2494787188 --outdir macs -n R1.chip

bamCompare -b1 R1.chip.dedup.sorted.gc.corrected.bam -b2
    ↪ R1.input.dedup.sorted.gc.corrected.bam --scaleFactorsMethod None
    ↪ --effectiveGenomeSize 2494787188 -p 10 --operation log2 --normalizeUsing RPKM -bl
    ↪ blacklist.bed -o R1.chip.input.log2.bw

computeMatrix reference-point -a 2000 -b 2000 --referencePoint center --smartLabels -R
    ↪ peaks.bed -S R1.chip.input.log2.bw -bs 5 -p 48
```

```
meme-chip -o R1.chip.meme -neg random.ATAC.fasta -order 2 -meme-p 12 -meme-nmotifs 40
    ↪ -psp-gen R1.chip.peaks.fasta
```

##### Command-line arguments for RNA-seq analysis

```
sailfish quant -l IU -i gencode.M23.index -1 R1.1.fq -2 R1.2.fq --biasCorrect -g gencode.M23.genes
    ↪ --numBootstraps 20 -o outdir -p 12

bedtools makewindows -g GRCm38.p6.fa.fai -w 1000 -i srcwinnum > GRCm38.p6.1kb.bed

bedtools nuc -fi GRCm38.p6.fa -bed GRCm38.p6.1kb.bed > GRCm38.p6.1kb.nuc

computeMatrix scale-regions -m 10000 -a 2000 -b 5000 -R gencode.M23.genes.bed -S GRCm38.p6.AT.bw -out
    ↪ gencode.M23.genes.AT.matrix.gz
```

##### R Script for differential gene expression of *Sall4* mutants

```
library(BiocParallel)
library(DESeq2)
register(MulticoreParam(4))

deseq_function <- function(counts_file, design_file, threshold, out_prefix){
  counts = read.csv(counts_file, sep="\t", header = TRUE,
                    row.names = 1, check.names = FALSE)
  design = read.csv(design_file, header=TRUE, sep=",", row.names=1)

  dds <- DESeqDataSetFromMatrix(countData = counts,
                                colData = design,
                                design = ~ condition)

  dds <- dds[rowSums(counts(dds)) > threshold,]

  # Performing DESeq2 analysis
  dds <- DESeq(dds, parallel=TRUE)
  saveRDS(dds, file=paste(out_prefix, "dds.rds", collapse="", sep=""))
  rld <- rlog(dds)

  ko_vs_wt <- results(dds, c("condition", "KO", "WT"), independentFiltering = TRUE)
  write.table(as.data.frame(ko_vs_wt),
      file=paste( out_prefix, "ko_vs_wt.tsv", collapse = "", sep=""),
      quote=F, col.names=NA, sep="\t")
  print(paste(c(counts_file, "finished")))
}
```

**R Script for analysing genotype-specific differences over time during stem cell differentiation**

```
library(BiocParallel)
library(DESeq2)
register(MulticoreParam(4))

find_hull <- function(df) df[chull(df$PC1, df$PC2), ]

deseq_function <- function(counts_file, design_file, out_prefix){
  counts = read.csv(counts_file, sep="\t", header = TRUE,
                    row.names = 1, check.names = FALSE)
  design = read.csv(design_file, header=TRUE, sep="\t", row.names=1)
  design$name <- relevel(design$name, "WT")
  design$timepoint <- as.factor(design$timepoint)

  dds <- DESeqDataSetFromMatrix(countData = counts,
                                colData = design,
                                design = ~ name + timepoint + name:timepoint)

  # Performing DESeq2 analysis
  dds <- DESeq(dds, parallel=TRUE)
  saveRDS(dds, file=paste(out_prefix, "dds.rds", collapse="", sep=""))
  rld <- rlog(dds)

  ddsTC <- DESeq(dds, test="LRT", reduced = ~ name + timepoint)
  resTC <- results(ddsTC)

  write.table(assay(rld), file=paste(c(out_prefix, "rld.tsv"), collapse="", sep=""), sep="\t")
  write.table(as.data.frame(resTC), file=paste(c(out_prefix, "fc.tsv"), collapse="", sep=""), sep="\t")

  print(paste(c(counts_file, "finished")))
}
```

**Linear Regression Model**

```
import pandas as pd
import statsmodels.api as sm
from statsmodels.stats import multitest

# Fitting OLS linear regression model to data
df = pd.read_csv("fold_change_AT.tsv", sep="\t")
X = sm.add_constant(df[["AT mean"]])
y = df["log2FoldChange"].values
model = sm.OLS(y, X).fit()
at_hi_conf, at_low_conf = tuple(model.conf_int(0.01).loc["AT mean"].T.values)
at_mean = model.params.loc["AT mean"]
r_squared = model.rsquared
f_pvalue = model.f_pvalue

# Adjusting Type I errors
_, combined_df["FDR"], _, _ = multitest.multipletests(combined_df["f_pvalue"].values, alpha=0.01,
    ↪ method="fdr_bh")
```
